# Contributions of the four essential entry glycoproteins to HSV-1 tropism and the selection of entry routes

**DOI:** 10.1101/2020.03.10.985325

**Authors:** Adam T. Hilterbrand, Raecliffe E. Daly, Ekaterina E. Heldwein

## Abstract

Herpes Simplex viruses (HSV-1 and HSV-2) encode up to 16 envelope proteins, four of which are essential for entry. However, whether these four proteins alone are sufficient to dictate the broad cellular tropism of HSV-1 and the selection of different cell-type dependent entry routes is unknown. To begin addressing this, we previously pseudotyped VSV, lacking its native glycoprotein G, with only the four essential entry glycoproteins of HSV-1: gB, gH, gL, and gD. This novel VSVΔG-BHLD pseudotype recapitulated several important features of HSV-1 entry: the requirement for gB, gH, gL, gD, a cellular receptor, and sensitivity to anti-gB and anti-gH/gL neutralizing antibodies. However, due to the use of a single cell type in that study, the tropism of the VSVΔG-BHLD pseudotype was not investigated. Here, we show that the cellular tropism of the pseudotype is severely limited compared to wild-type HSV-1 and that its entry pathways differ from the native HSV-1 entry pathways. To test the hypothesis that other HSV-1 envelope proteins may contribute to HSV-1 tropism, we generated a derivative pseudotype containing the HSV-1 glycoprotein gC (VSVΔG-BHLD-gC) and observed a gC-dependent increase in entry efficiency in two cell types. We propose that the pseudotyping platform developed here has the potential to uncover functional contributions of HSV-1 envelope proteins to entry in a gain-of-function manner.

**Importance:** Herpes simplex viruses (HSV-1 and HSV-2) contain up to 16 different proteins in their envelopes. Four of these, glycoproteins gB, gD, gH, and gL, are termed essential with regard to entry whereas the rest are typically referred to as non-essential based on the entry phenotypes of the respective single genetic deletions. However, the single-gene deletion approach, which relies on robust loss-of-function phenotypes, may be confounded by functional redundancies among the many HSV-1 envelope proteins. We have developed a pseudotyping platform, in which the essential four entry glycoproteins are isolated from the rest, which can be added back individually for systematic gain-of-function entry experiments. Here, we show the utility of this platform for dissecting the contributions of HSV envelope proteins, both the essential four and the remaining dozen (using gC as an example), to HSV entry.

## Introduction

Herpes simplex viruses (HSV-1 and HSV-2) are enveloped viruses that infect much of the world’s population for life and cause diseases ranging from painful oral or genital lesions to serious conditions such as encephalitis and blindness (1, 2). These viruses enter target cells by different, cell-type specific routes. For example, they enter neurons by direct fusion of their envelopes with the plasma membrane (3) and epithelial cells by endocytosis followed by fusion with an endosomal membrane (4, 5). While these entry pathways have been broadly described, the underlying mechanisms and the contributions of individual viral and cellular proteins to the selection of these entry routes remain incomplete.

HSV-1 entry by any route requires the coordinated efforts of four glycoproteins – gB, gH, gL, and gD, which are essential for entry (6–8) – and a cellular gD receptor (3, 9, 10). gB, gH, gL, and gD are also sufficient for cell-cell fusion of uninfected receptor-bearing cells expressing these four glycoproteins (11, 12). The prevalent model, which was largely developed through studies using the cell-cell fusion system, posits that these four viral glycoproteins orchestrate membrane fusion through a sequential activation process termed a cascade (13, 14). First, gD binds one of its three cellular receptors, nectin-1, herpesvirus entry mediator (HVEM), or 3-O-sulfated heparan sulfate (3-OS-HS) (15). Binding of gD to its receptor triggers a conformational change within gD (14, 16, 17) that enables it to bind (18) and activate the gH/gL heterodimer (13, 19, 20). In turn, gH/gL presumably interacts with and activates gB (13, 21, 22), the fusogen that mediates the merger of the HSV lipid envelope with the cellular membrane (9, 10, 14, 23).

In addition to the essential four glycoproteins, HSV-1 encodes up to 12 more envelope proteins, eight glycosylated and four unglycosylated (24–26). Current models of HSV-1 entry do not account for the potential effects of these envelope proteins. Therefore, being able to functionally uncouple the four essential glycoproteins – gB, gH, gL, and gD – from the rest is fundamental for elucidating their contributions to HSV-1 cellular tropism and entry pathways.

One powerful system that enables such studies is the vesicular stomatitis virus (VSV)- based pseudotype, in which the native VSV glycoprotein, G, is replaced with a viral envelope protein of interest (27). The VSV pseudotyping system allows one to define entry mechanisms conferred by a specific viral glycoprotein by effectively isolating it from its native viral context. This platform has been used to elucidate the entry mechanisms of many viruses, notably those that require BSL-3 or BSL-4 containment facilities, including SARS-CoV (28, 29), SARS-CoV-2 (30), Ebola virus (31), Lassa virus (32, 33), Lujo virus (34), Hantavirus (35), Rift Valley fever virus (36), or those that are difficult to culture, such as Hepatitis C virus (37) or Japanese encephalitis virus (38). The use of VSV pseudotypes has been particularly useful in identifying cellular receptors of many viruses (31–36).

To determine whether the essential four HSV-1 glycoproteins were sufficient for entry, we previously generated VSV lacking its native glycoprotein G and pseudotyped with HSV-1 gB, gH, gL, and gD through *trans*-complementation (VSVΔG-BHLD) (39). The VSVΔG-BHLD pseudotype efficiently entered C10 cells (B78 murine melanoma cells expressing HSV-1 receptor nectin-1), and its entry – like that of HSV-1 – required gB, gH, gL, gD, and a gD receptor, and was inhibited by anti-gB and anti-gH/gL neutralizing antibodies (39). However, this study left unknown whether the VSVΔG-BHLD pseudotype could enter any HSV-1 susceptible cell types or utilize native entry routes. Therefore, we sought to directly compare the cellular tropism and the entry pathways of the VSVΔG-BHLD pseudotype and HSV-1 to determine the extent by which these were conferred solely by the four essential glycoproteins.

Here, we expanded our studies to six additional HSV-1 susceptible cell lines. VSVΔG-BHLD was only able to enter, with reasonable efficiency, two out of the seven HSV-1-susceptible cell lines. Additionally, the VSVΔG-BHLD pseudotype entered both cell lines by routes different from those used by HSV-1. Differences in tropism and routes of entry could not be accounted for by either cell-surface receptor levels, their nature (nectin-1 vs. HVEM), the relative amounts of gB, gH, gL, and gD, or virion morphology (VSV vs. HSV-1). Therefore, we conclude that the four essential HSV-1 entry glycoproteins are insufficient for entry into any HSV-1-susceptible cell and do not specify native entry routes. Our results raise an intriguing possibility that HSV-1-specific components outside the essential four glycoproteins influence HSV-1 entry. Indeed, when the HSV-1 glycoprotein gC was included in the VSVΔG-BHLD pseudotype (VSVΔG-BHLD-gC), entry efficiency into CHO-HVEM and HaCaT cells increased, suggesting a cell-type dependent gain-of-function conferred by gC. Therefore, we hypothesize that the so-called non-essential HSV-1 envelope proteins, which are missing from the VSVΔG-BHLD pseudotype, are important in specifying both HSV-1 tropism and its routes of cell entry.

## Results

### VSVΔG-BHLD pseudotypes enter a limited repertoire of HSV-1 susceptible cells

To determine the tropism of the VSVΔG-BHLD pseudotype, we selected seven HSV-1-susceptible cell lines (Fig. 1). The cell lines B78H1 and CHO-K1, which lack HSV-1 receptors served as negative controls. HSV-1 efficiently infected all seven receptor-bearing cells but not the receptor-negative cells (Fig. 1A). The VSVΔG-BHLD pseudotype efficiently infected C10 cells (Fig. 1B), consistent with our previous report (39). The VSVΔG-BHLD pseudotype also infected CHO-HVEM, CHO-nectin-1, and HaCaT cells, albeit with lower efficiency (Fig. 1B). Although VSVΔG-BHLD entry into CHO-HVEM and CHO-nectin-1 cells was relatively inefficient, it was clearly receptor-dependent (Fig. 1B). However, no measurable VSVΔG-BHLD entry was observed in HeLa, Vero, or SH-SY5Y cells (Fig. 1B) even at MOI of 10 (Fig. S1A).

**Fig. 1.**
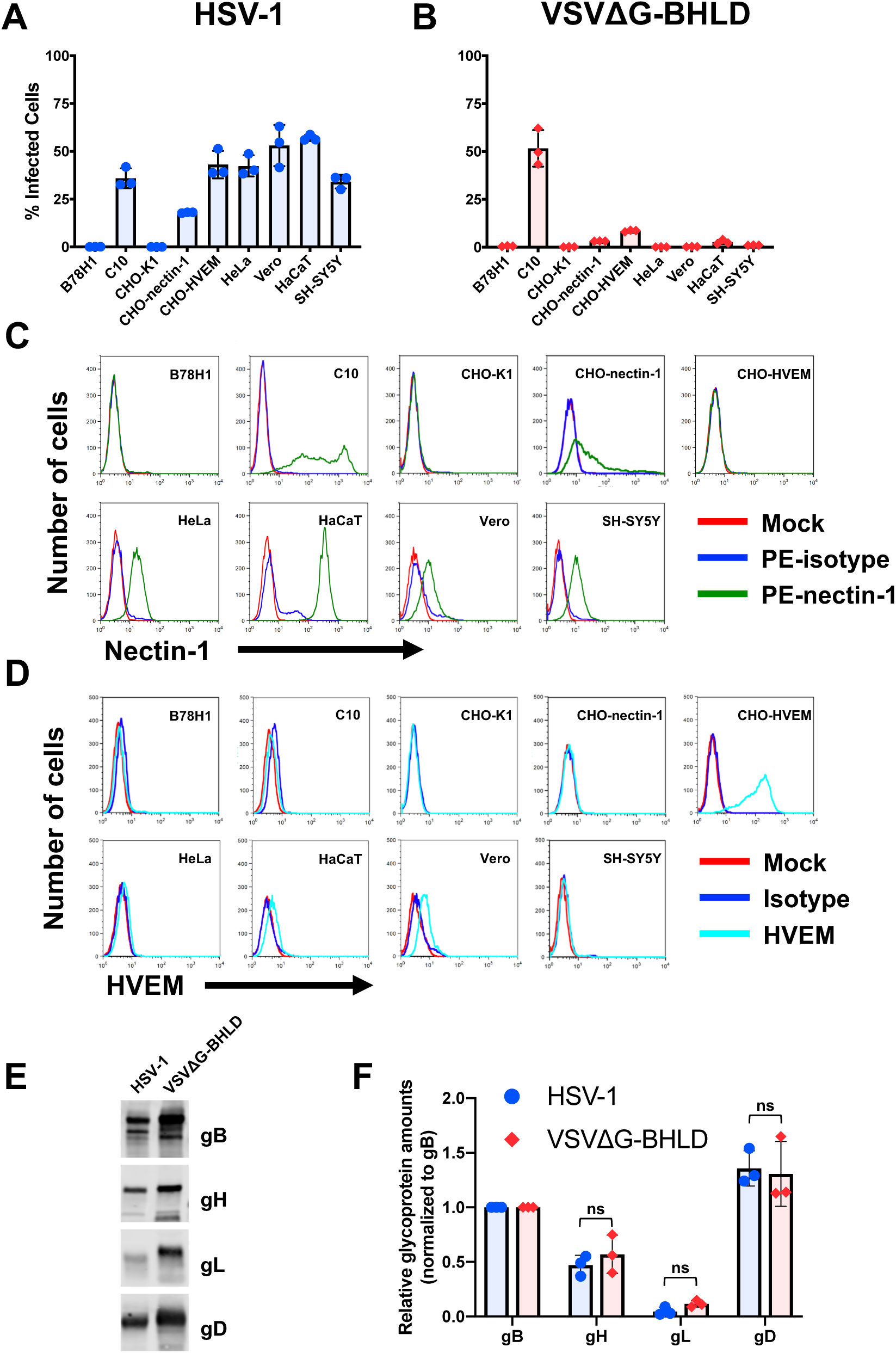
VSVΔG-BHLD pseudotype narrow cellular tropism is not due to differences in relative glycoprotein ratios or receptor expression levels. HSV-1 (A) and VSVΔG-BHLD (B) entry was assessed on nine cell lines, B78H1, C10, CHO-K1, CHO-nectin-1, CHO-HVEM, HeLa, Vero, HaCaT, and SH-SY5Y. Cells were infected at a MOI of 1. Entry was quantitated by flow cytometry at 6 hours post infection. C) Cell surface expression of nectin-1 (C) and HVEM (D) was analyzed by flow cytometry. Surface levels of nectin-1 were quantitated by staining cells with an anti-nectin-1 monoclonal antibody CK41 conjugated to phycoerythrin (PE) (green histograms). Surface levels of HVEM were quantitated by staining cells with an anti-HVEM polyclonal antibody R140 and a FITC-labeled secondary antibody (cyan). Blue histograms are isotype controls. Red histograms are mock (no antibody) controls. E and F) Relative ratios of gB:gH:gL:gD for HSV-1 and VSVΔG-BHLD particles. HSV-1 and VSVΔG-BHLD virions were purified either through a continuous sucrose gradient (HSV-1) or continuous Optiprep gradient (VSVΔG-BHLD), pelleted, and analyzed for their glycoprotein content (gB, gH, gL, and gD) by western blot (anti-gB pAb R68, anti-gH pAb R137, anti-gL mAb L1, and anti-gD pAb R7). A representative western blot is shown. The amounts of gB, gH, gL, and gD in three different virion preparations were determined by densitometry. Levels of gH, gL, and gD were normalized to gB levels from their respective virions. A Student’s T-test with Welch’s correction was used to determine the significance of differences between relative amounts of gH, gL, or gD between HSV-1 and VSVΔG-BHLD virions (ns = not significant; p < 0.05 = *; p < 0.01 = **; p < 0.001 = ***).

Two additional VSV pseudotypes were used as controls. VSVΔG-G is VSVΔG pseudotyped *in trans* with native VSV glycoprotein G (39). VSVΔG-PIV5 is VSVΔG pseudotyped with entry glycoproteins HN and F from parainfluenza virus 5 (PIV5) (40). Both controls infected all 9 tested cell lines, with varying efficiency (Figs. S1B and C), suggesting the limited tropism of VSVΔG-BHLD could not be attributed to VSV morphology alone.

### Cell-surface receptor levels do not correlate with differences in VSVΔG-BHLD cellular tropism

We first asked whether differences in cell surface levels of HSV-1 receptors could account for VSVΔG-BHLD entry efficiency. Levels of HSV-1 gD receptors nectin-1 and HVEM vary across cell lines, and susceptibility to HSV-1 infection generally correlates with surface receptor levels (41). Surface levels of nectin-1 and HVEM were measured in all 9 cell lines by flow cytometry (Figs. 1C and D). As expected, neither receptor was detected on the receptor-negative cell lines B78H1 and CHO-K1 (Figs. 1C and D). C10 and HaCaT cells had the highest levels of nectin-1, whereas intermediate levels of nectin-1 were detected on CHO-nectin-1, HeLa, Vero and SH-SY5Y cells (Fig. 1C). CHO-HVEM cells had high levels of HVEM but no detectable nectin-1 on their surface (Fig. 1D). In addition to nectin-1, we also detected HVEM on the surface of HaCaT and Vero cells (Fig. 1D), but the low amounts of HVEM suggested that nectin-1 likely functions as the primary receptor in these cells. Surprisingly, while both C10 and HaCaT cells expressed high levels of nectin-1, the VSVΔG-BHLD pseudotype efficiently entered only C10 cells. These results suggest that surface receptor levels alone do not explain the varying entry efficiencies of the VSVΔG-BHLD pseudotype into the tested cell lines.

### Differences in tropism of VSVΔG-BHLD and HSV-1 do not correlate with the relative gB:gH:gL:gD ratios

HSV-1 and VSV acquire their envelopes from different sources: Trans Golgi Network (TGN) or endosomes for HSV-1 (42, 43) vs. the plasma membrane (PM) for VSV (27). Different envelope origins could affect the gB:gH:gL:gD ratios on viral particles and, possibly, influence entry efficiency. To test this hypothesis, purified HSV-1 and VSVΔG-BHLD particles were analyzed for gB, gH, gL, and gD content by western blot (Fig. 1E), and relative gB:gH:gL:gD ratios were determined by densitometry. In each virus, levels of gH, gL, and gD were normalized to their respective gB levels. We found that the gB:gH:gL:gD ratios in HSV-1 (1:0.47:0.05:1.36) and VSVΔG-BHLD (1:0.57:0.12:1.31) virions were similar (Fig. 1F) and unlikely to account for the observed differences in tropism.

### Entry of both VSVΔG-BHLD and HSV-1 into C10 and CHO-HVEM cells occurs by endocytosis

HSV-1 can enter different cell types by fusion at the plasma membrane (Vero and SH-SY5Y) (5, 44) or by endocytosis (C10, CHO-nectin-1, CHO-HVEM, HeLa, and HaCaT) (3, 5, 45, 46). To compare the entry routes of the VSVΔG-BHLD pseudotype and HSV-1, we chose C10 and CHO-HVEM cells because the VSVΔG-BHLD pseudotype infected C10 (∼50%) or CHO-HVEM (∼8%) cells to an appreciable extent and in a receptor-dependent manner (Fig. 1B).

We first treated cells with a hypertonic solution of sucrose, a broad inhibitor of endocytic pathways (47, 48). Entry of both HSV-1 and VSVΔG-BHLD into C10 and CHO-HVEM cells was inhibited by sucrose (Fig. 2A-D), implicating endocytosis. As a control, sucrose also prevented the endocytic uptake of Alexa Fluor 488-labeled transferrin into both C10 and CHO-HVEM cells (Fig. S2E). As expected, sucrose did not inhibit entry of VSVΔG-PIV5 into either cell line (Figs. S2B and D) because PIV5 enters by fusion at the plasma membrane (40, 49). VSV entry occurs by endocytosis (50–52), and, accordingly, sucrose blocked entry of VSVΔG-G into C10 cells (Fig. S2A). Surprisingly, it did not block entry into CHO-HVEM cells (Fig. S2C), suggesting that the inhibitory effect of hypertonic sucrose on VSV-G-dependent entry is cell-type specific.

**Fig. 2.**
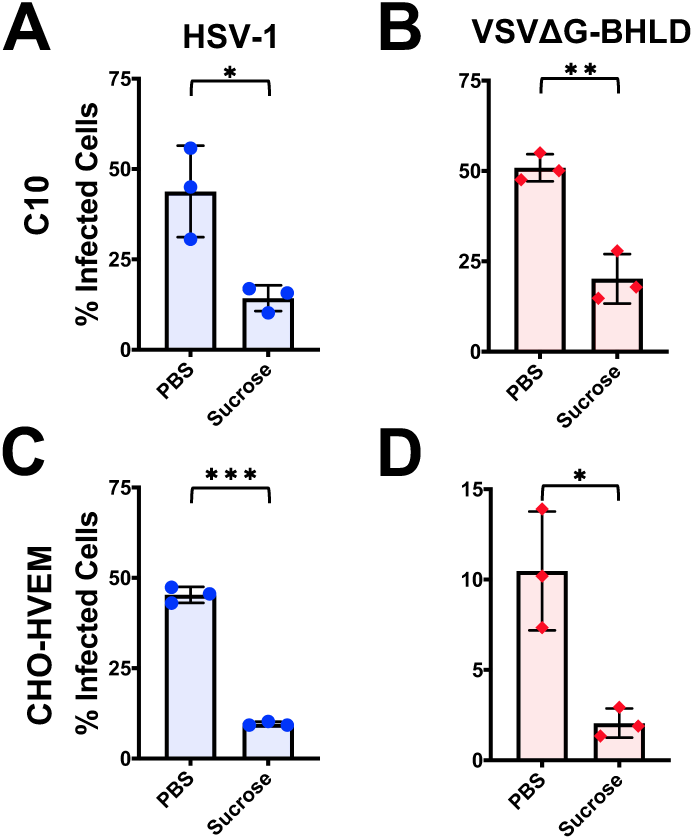
HSV-1 and VSVΔG-BHLD enter cells by endocytosis. C10 (A and B) and CHO-HVEM (C and D) cells were pretreated with a hypertonic solution of sucrose (0.3 M) and infected with HSV-1 and VSVΔG-BHLD at MOI = 1. Infectivity was quantitated by flow cytometry at 6 hours post infection. Significance was calculated using a two-tailed Student’s T-test with Welch’s correction (p < 0.05 = *; p < 0.01 = **; p < 0.001 = ***).

### VSVΔG-BHLD entry requires dynamin but not clathrin whereas HSV-1 entry requires both

Having established that the VSVΔG-BHLD pseudotype entered cells by endocytosis, we next sought to identify the entry routes and compare them to those of HSV-1 by using both chemical and genetic means of inhibiting various endocytic uptake pathways.

First, we examined the role of clathrin-mediated endocytosis (CME), one of the most well studied endocytic pathways hijacked by viruses for entry (53, 54). CME requires both clathrin, to promote receptor-mediated endocytosis (55), and dynamin, a GTPase that mediates scission of the endocytic vesicle (56). We chose three commonly used dynamin inhibitors: Dynasore, Dyngo-4a, and myristyltrimethylammonium bromide (MiTMAB) (47, 57). For clathrin inhibition, we chose Pitstop-2, which selectively blocks CME by preventing ligand association with the clathrin terminal domain (47). Another CME inhibitor, chlorpromazine, which prevents clathrin association with the plasma membrane (47), was also tested but found to be toxic to both C10 and CHO-HVEM cells. Inhibitory activity of all four compounds was ascertained by their ability to inhibit CME of transferrin (Figs. S3E and F).

HSV-1 entry into both C10 and CHO-HVEM cell lines was inhibited by all four inhibitors (Figs. 3A and B) indicating that HSV-1 enters both cell lines by CME. Entry of the VSVΔG-BHLD pseudotype into C10 and CHO-HVEM cells was sensitive to all dynamin inhibitors (Figs. 3C and D). However, Pitstop-2 did not block VSVΔG-BHLD entry (Figs. 3C and D), suggesting that CME was not involved in entry. Collectively, these observations suggested that HSV-1 enters both C10 and CHO-HVEM cells by CME whereas VSVΔG-BHLD pseudotypes utilize a dynamin-dependent, clathrin-independent entry route. These results were the first indication of possible differences in entry routes of HSV-1 and VSVΔG-BHLD.

**Fig. 3.**
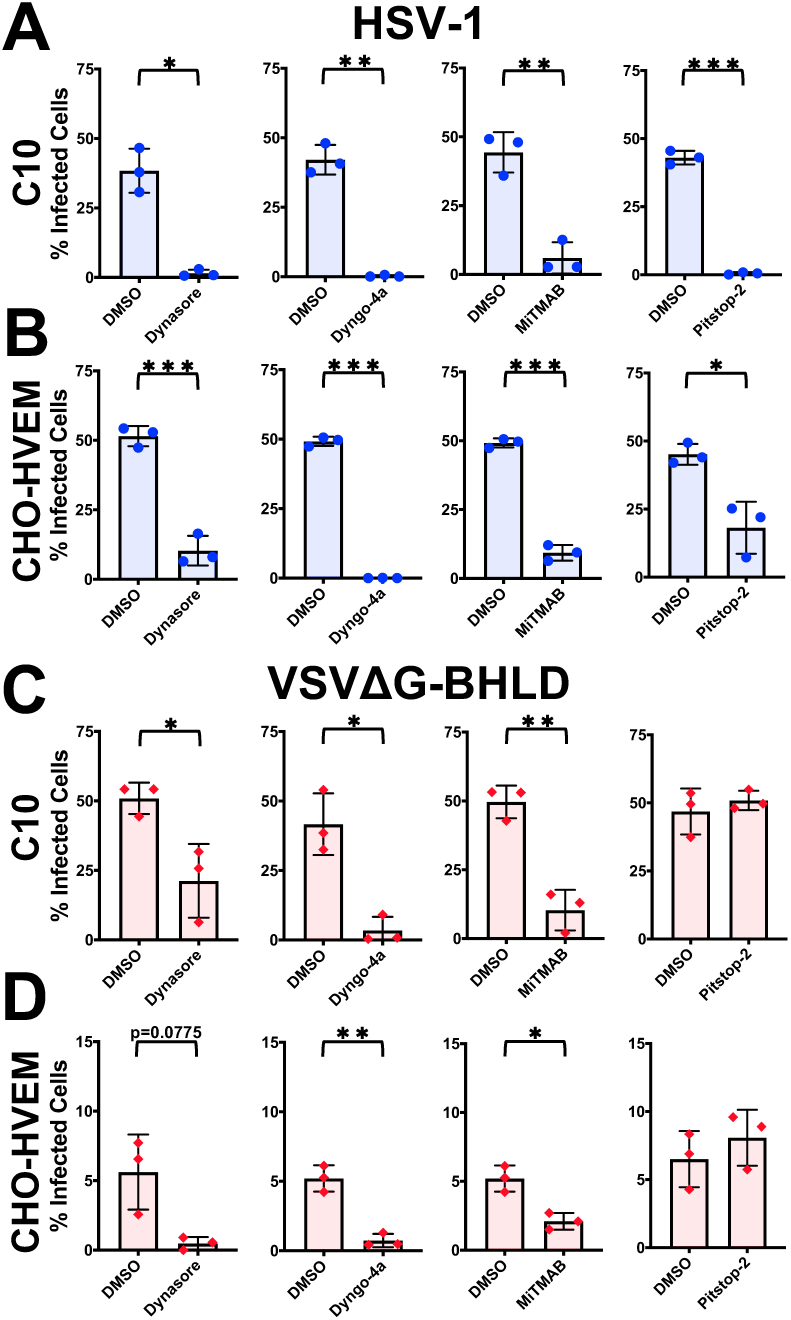
Both HSV-1 and VSVΔG-BHLD require dynamin but only HSV-1 requires clathrin for entry. C10 (A and C) and CHO-HVEM (B and D) cells were pretreated with dynamin inhibitors Dynasore (80 μM), Dyngo-4a (25 μM), MiTMAB (5 μM), or the CME inhibitor, Pitstop-2 (30 μM) and infected with HSV-1 or VSVΔG-BHLD at MOI = 1. Infectivity was quantitated by flow cytometry at 6 hours post infection. CHO-HVEM cells treated with Dyngo-4a or MiTMAB used the same DMSO control, as indicated by the same bar graph appearing twice each in panels C and D. Significance was calculated using a two-tailed Student’s T-test with Welch’s correction (p < 0.05 = *; p < 0.01 = **; p < 0.001 = ***).

Entry of the control VSVΔG-G pseudotype into C10 cells was blocked by all three dynamin inhibitors and the clathrin inhibitor (Fig. S3A), and its entry into CHO-HVEM cells was blocked by Pitstop-2 (Fig. S3C) and by two of three dynamin inhibitors, Dynasore and Dyngo-4a (Fig. S4C), strongly implicating CME as the entry route (52, 58). VSVΔG-PIV5 entry into CHO-HVEM cells was not blocked by any of the four inhibitors (Fig. S3D) while its entry into C10 cells was blocked only by one out of three dynamin inhibitors, Dyngo-4a (Fig. S3C), consistent with the previous report of entry into other cell types by fusion at the plasma membrane (40, 49).

### Cholesterol is important for entry of both VSVΔG-BHLD and HSV-1

Our results suggested that VSVΔG-BHLD did not use CME for entry into either C10 or CHO-HVEM cells, implicating a clathrin-independent endocytic (CIE) route. Caveolin-dependent endocytosis is a major CIE (59) that is hijacked by viruses such as SV40 or Japanese encephalitis virus (60, 61). Caveolin-1 is cellular protein that, similarly to clathrin, promotes membrane curvature and subsequent endocytosis through the formation of caveolae (62). Caveolin-dependent entry requires plasma membrane cholesterol for proper caveolin-1 association with the membrane (62, 63).

Previous work has demonstrated that cellular cholesterol was important for HSV-1 entry into C10 cells (64). Similarly, entry of both HSV-1 and VSVΔG-BHLD into C10 and CHO-HVEM cells decreased when cholesterol was removed from cellular membranes using a cholesterol-depleting agent, methyl-β-cyclodextrin (MβCD) (Figs. 4A-D). Efficiency of cholesterol depletion was confirmed by the reduction of the cholesterol-dependent association of cholera toxin subunit B (CTB) (65) with C10 and CHO-HVEM cells upon MβCD treatment (Fig. S4E). Entry of VSVΔG-G was insensitive to cholesterol depletion (Figs. S4A and C), as reported for other cell types (66, 67). In the case of VSVΔG-PIV5, only entry into C10 cells was cholesterol-dependent (Fig. S4B) whereas the entry into CHO-HVEM cells was not (Fig. S4D), suggesting that cholesterol is required for VSVΔG-PIV5 entry in a cell-type dependent manner.

**Fig. 4.**
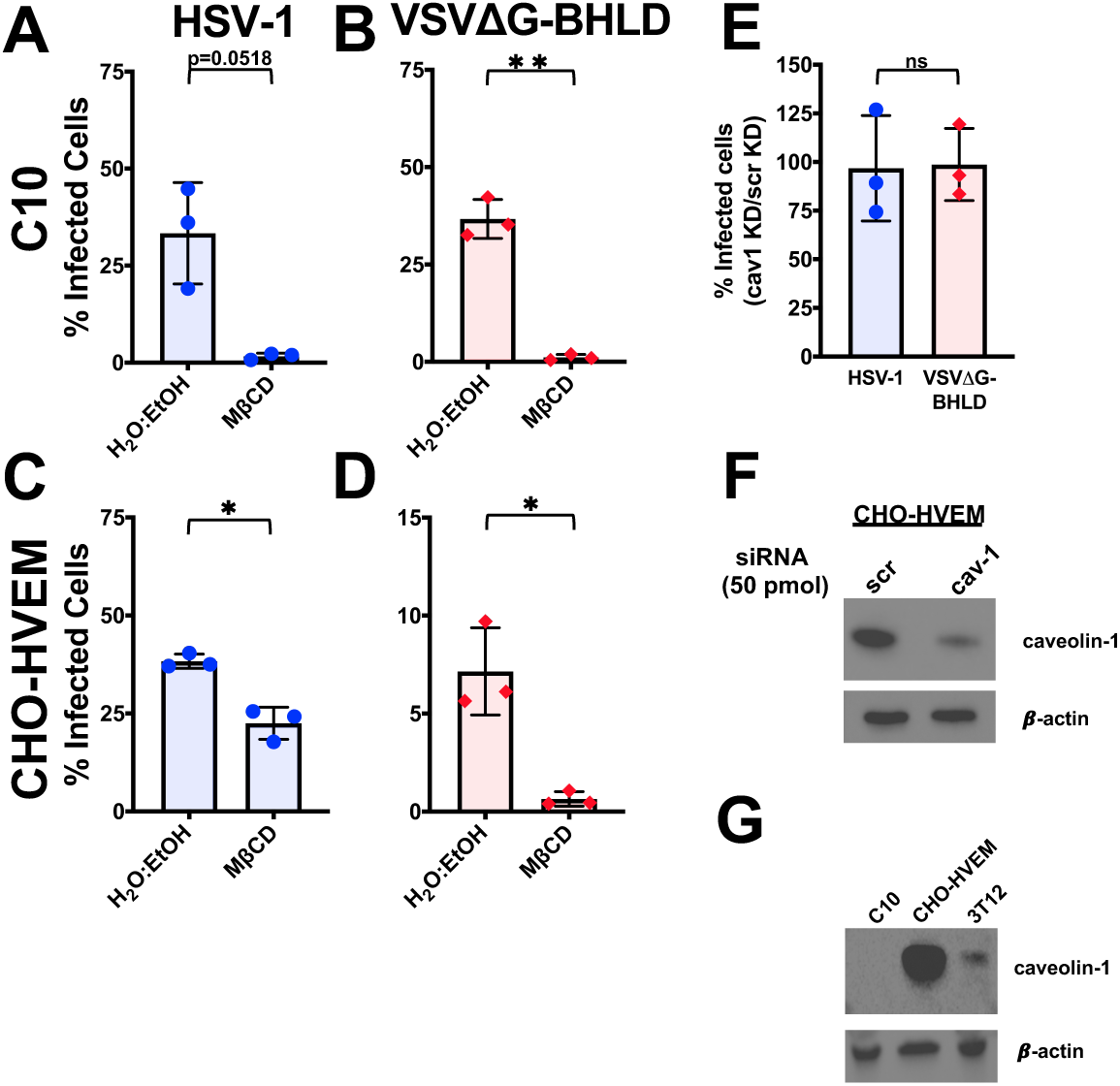
HSV-1 and VSVΔG-BHLD entry requires cellular cholesterol but not caveolin-1. C10 (A and B) and CHO-HVEM (C and D) cells were pretreated with a cholesterol-removal drug methyl-β-cyclodextran, MβCD (5 mM) and infected with HSV-1 (A and C) or VSVΔG-BHLD (B and D) at MOI = 1. Infectivity was quantitated by flow cytometry at 6 hours post infection. E) CHO-HVEM cells were transfected with a caveolin-1 siRNA (cav-1) or a scrambled control siRNA (scr) (both 50 pm) and infected with HSV-1 or VSVΔG-BHLD at MOI = 1. Infectivity was quantitated by flow cytometry at 6 hours post infection. Significance was calculated using a two-tailed Student’s T-test with Welch’s correction (p < 0.05 = *; p < 0.01 = **; p < 0.001 = ***). F) Western blot analyses, using antibody clone 4H312 (Santa Cruz Biotechnology), of caveolin-1 knockdown in CHO-HVEM cells (representative of three western blots, one from each biological replicate). G) Western blot analyses, using antibody clone 4H312 (Santa Cruz Biotechnology), of caveolin-1 levels in C10, CHO-HVEM, and 3T12 cells.

Entry of neither HSV-1 nor VSVΔG-BHLD into CHO-HVEM cells was reduced by the knockdown of caveolin-1 (Fig. 4E), similarly to the VSVΔG-G and VSVΔG-PIV5 control viruses (Fig. S4F). Successful knockdown was verified by western blot (Fig. 4F). Surprisingly, no caveolin-1 was detected in C10 cells (Fig. 4G). Caveolin-1 was detected in 3T12 cells (murine fibroblasts), which ruled out species-dependent recognition of murine vs. hamster caveolin-1, (Fig. 4G). Therefore, C10 cells appear to express no detectable caveolin-1. We conclude that while cellular cholesterol is important for the entry of both HSV-1 and VSVΔG-BHLD, neither virus utilizes caveolin-1-mediated endocytosis for entry into C10 and CHO-HVEM cells.

### HSV-1 and VSVΔG-BHLD do not enter C10 and CHO-HVEM cells by macropinocytosis, but NHE1 and Rac1 are important for optimal VSVΔG-BHLD entry

We next evaluated the potential involvement of macropinocytosis, another CIE commonly used by viruses (68). We selected three known inhibitors of macropinocytosis: cytochalasin D (CytoD), 5-(N-Ethyl-N-isopropyl)amiloride (EIPA), and NSC23766 (47). CytoD is a potent inhibitor of actin polymerization that disrupts filamentous actin (47). EIPA blocks macropinocytosis by blocking Na^+^/H^+^ exchange proteins (NHE), which decreases the intracellular pH and inhibits small GTPase function important for macropinocytosis (69). NSC23766 blocks macropinocytosis by inhibiting the activity of the small GTPase Rac1 (70). Inhibitory activity of all three compounds was confirmed by their ability to inhibit macropinocytosis of rhodamine-B-labeled 70-kDa dextran (Fig. S7E).

HSV-1 entry into either C10 or CHO-HVEM cells was not appreciably inhibited by any of the macropinocytosis inhibitors (Figs. 5A and C). By contrast, VSVΔG-BHLD entry into both C10 and CHO-HVEM cells was reduced by NSC23766 and, to a lesser extent, EIPA but not by cytochalasin D (Figs. 5B and D). While the inhibitory effect of NSC23766 and EIPA would appear to implicate macropinocytosis as the route of VSVΔG-BHLD entry into C10 and CHO-HVEM cells, the lack of actin involvement argues against it. This is because assembly of filamentous actin is essential for the formation of membrane ruffles and subsequent uptake during macropinocytosis (71). Actin polymerization is, thus, a major hallmark of macropinocytosis (68). Accordingly, we did not observe any appreciable co-localization of VSVΔG-BHLD virions with the 70-kDa rhodamine B labeled dextran, a fluid phase uptake marker (Figs. S5 and S6). Collectively, these results suggest that neither VSVΔG-BHLD nor HSV-1 utilize macropinocytosis for entry into C10 or CHO-HVEM cells. However, NHE and Rac1 facilitate VSVΔG-BHLD entry into both cell lines, presumably, independently of their role in macropinocytosis.

**Fig. 5.**
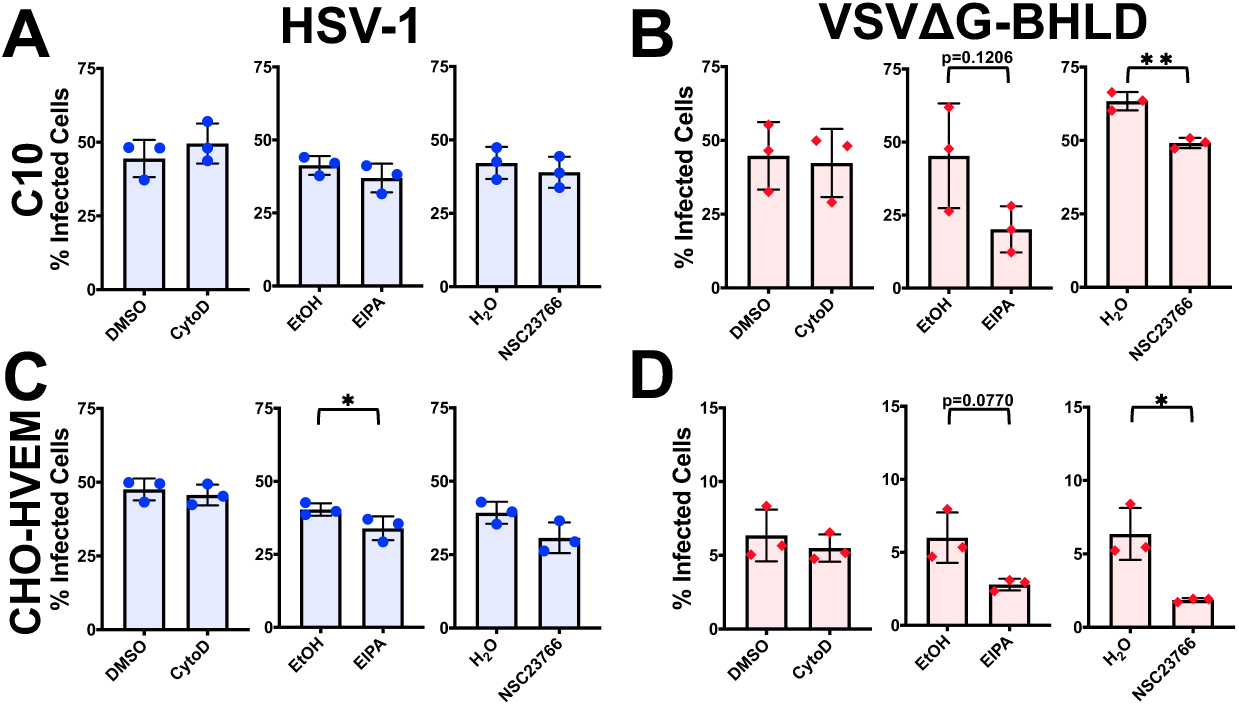
Neither HSV-1 nor VSVΔG-BHLD entry requires macropinocytosis. C10 (A and B) and CHO-HVEM (C and D) cells were pretreated with macropinocytosis inhibitors cytochalasin D (2 μM), EIPA (25 μM), or NSC23766 (200 μM) and infected with HSV-1 or VSVΔG-BHLD at MOI = 1. Infectivity was quantitated by flow cytometry at 6 hours post infection. Significance was calculated using a two-tailed Student’s T-test with Welch’s correction (ns = not significant; p < 0.05 = *; p < 0.01 = **; p < 0.001 = ***).

Although VSV enters cells by CME rather than macropinocytosis, it has been shown to require filamentous actin to achieve full engulfment of the viral particle by the plasma membrane during endocytosis, as observed in BSC-1 cells, an African green monkey epithelial cell line (58, 72). Indeed, VSVΔG-G entry into C10 cells was modestly reduced by cytochalasin D and EIPA but not NSC23766 (Fig. S7A) whereas its entry into CHO-HVEM cells was somewhat reduced by EIPA but not cytochalasin D or NSC23766 (Fig. S7C). The apparent lack of actin involvement in VSVΔG-G entry into CHO-HVEM cells may represent a cell-type specific phenomenon. VSVΔG-PIV5 entry was not blocked with any of the three inhibitors of macropinocytosis, consistent with fusion at the plasma membrane, and was, in fact, increased in the presence of EIPA in C10 cells (Fig. S7B).

### HSV-1 and VSVΔG-BHLD differ in their requirements for Rab GTPases for entry

Endocytic entry by many viruses requires small GTPases known as Rabs. Rab GTPases are important for the formation of endosomal compartments in the cell (73). Different viruses penetrate endocytic membranes at distinct endosomal maturation stages – for example, VSV fuses with membranes of early endosomes whereas influenza A virus fuses with membranes of late endosomes (74) – so, proper formation of these endosomal compartments is essential for viral entry. GTPases Rab5 and Rab7 are important for the maturation and formation of early endosomes and late endosomes/multi-vesicular bodies (MVBs), respectively (75). Overexpression of dominant negative (DN) forms of either Rab5 (Rab5DN) or Rab7 (Rab7DN) suppress early and late endosome formation, respectively (76, 77).

To identify the endosomal compartment(s) required for HSV-1 or VSVΔG-BHLD entry, C10 and CHO-HVEM cells were transfected with constructs encoding fluorescently tagged Rab5DN or Rab7DN and then infected. The transfection efficiency was 40-50% in C10 cells and 30-40% in CHO-HVEM cells as measured by flow cytometry. To determine entry efficiency by flow cytometry, viral entry was calculated by dividing the percent of infected and transfected cells by the total number of transfected cells. Overexpression of either Rab5DN or Rab7DN had no significant effect on HSV-1 entry into either cell line (Figs. 6A and E). These results in C10 and CHO-HVEM cells agree with recent work that indicates HSV-1 enters CHO-HVEM cells by a non-canonical endocytic route, independent of Rab5 or Rab7 (78). VSVΔG-BHLD entry into CHO-HVEM cells was also unaffected by either Rab5DN or Rab7DN (Fig. 6G). However, VSVΔG-BHLD entry into C10 cells was reduced in the presence of Rab5DN and, to some extent, Rab7DN (Fig. 6C). We hypothesize that HSV-1 entry into both cell lines and VSVΔG-BHLD entry into CHO-HVEM cells either do not depend on the endosomal maturation or occurs very early after internalization, prior to the formation of early endosomes. By contrast, VSVΔG-BHLD likely enters C10 cells out of early endosomes. This observation marked the first instance of a difference in VSVΔG-BHLD entry into C10 vs. CHO-HVEM cells.

**Fig 6:**
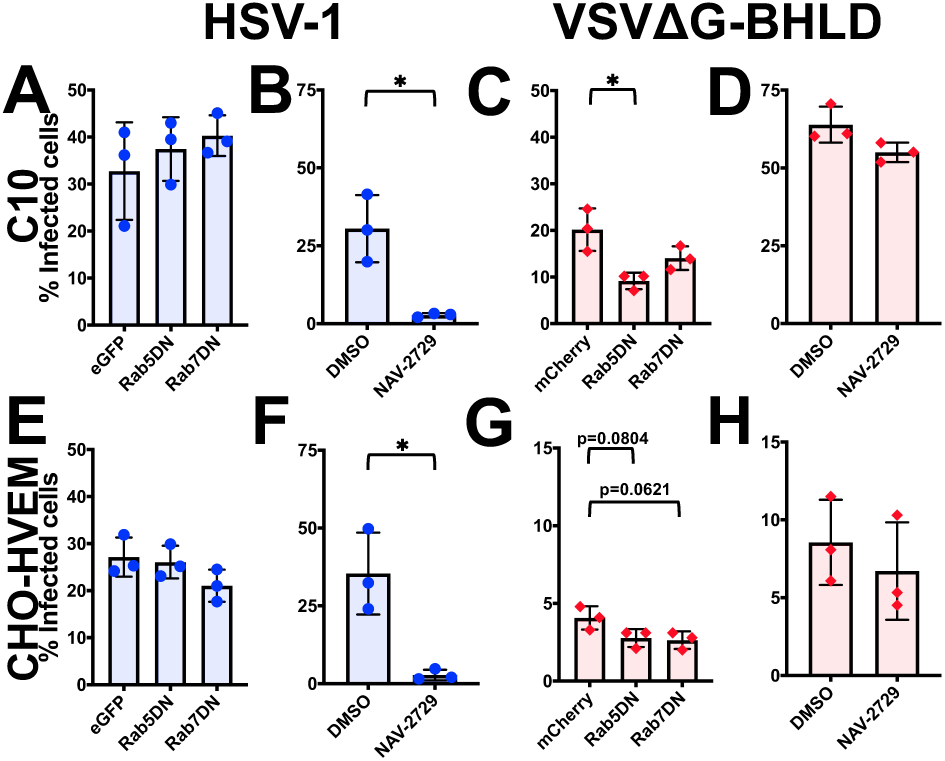
Roles of Rabs 5 and 7 and Arf6 in HSV-1 and VSVΔG-BHLD entry. The roles of the small GTPases Rab5, Rab7 (A, C, E, G), and Arf6 (B, D, F, H) were assessed for HSV-1 (A, B, E, F) and VSVΔG-BHLD (C, D, G, H) entry into C10 (A, B, C, D) and CHO-HVEM (E, F, G, H) cells. C10 (A and C) and CHO-HVEM (E and G) cells were transfected with either an empty vector control (eGFP or mCherry), eGFP or mCherry-tagged Rab5 dominant negative (DN), or eGFP or mCherry-tagged Rab7DN. Cells were infected at an MOI = 1 with either HSV-1 or VSVΔG-BHLD. Entry was assessed by flow cytometry at 6 hpi. The percent of infected cells was determined by dividing the number of virus(+)eGFP/mCherry(+) cells by the total number of eGFP/mCherry(+) cells. C10 (B and D) and CHO-HVEM cells (F and H) were treated with the Arf6 inhibitor NAV-2729 (25 μM) and infected with either HSV-1 or VSVΔG-BHLD at an MOI = 1. Significance was calculated using a two-tailed Student’s T-test with Welch’s correction (ns = not significant; p < 0.05 = *; p < 0.01 = **; p < 0.001 = ***).

As expected, entry of VSVΔG-G into both C10 and CHO-HVEM cells was reduced in the presence of Rab5DN (Figs. S8A and E), in accordance with reports of VSV fusing with membranes of early endosomes (52). VSVΔG-PIV5 entry into C10 cells was insensitive to Rab5DN or Rab7DN (Fig. S8C). Although VSVΔG-PIV5 entry into CHO-HVEM cells was reduced by Rab5DN and Rab7DN in a statistically significant manner, the differences were relatively small (Fig. S8G). This suggested that Rab5DN and Rab7DN have a minimal effect on VSVΔG-PIV5 entry as expected for a virus that fuses with the plasma membrane.

Another small GTPase, ADP-ribosylation factor 6 (Arf6), which is involved in regulating vesicular trafficking (79), regulates the endocytic entry of HIV, Coxsackievirus, and Vaccinia virus (80–82). To probe the role of Arf6 in entry, cells were treated with NAV-2729, which blocks Arf6 interaction with guanine exchange factors (GEFs) thereby preventing its activation (83). HSV-1 entry into both C10 and CHO-HVEM cells was inhibited by NAV-2729 (Figs. 6B and F) whereas VSVΔG-BHLD entry was not (Figs. 6D and H), which suggests that only HSV-1 entry requires Arf6 activity.

VSVΔG-PIV5 entry was not inhibited by NAV-2729 in either C10 or CHO-HVEM cells (Figs. S8D and H). However, VSVΔG-G entry was inhibited by NAV-2729 in both C10 and CHO-HVEM cells, suggesting that Arf6 could be involved in VSV endocytosis (Figs S8B and F). These results point to a previously unappreciated role of Arf6 in HSV-1 and VSV entry.

### Entry of the VSVΔG-BHLD pseudotype requires endosomal acidification in a cell-dependent manner

To investigate the role of endosomal acidification in entry, we used three common inhibitors, NH_4_Cl, a weak base; monensin, a carboxylic ionophore; and bafilomycin A1 (BFLA), an endosomal V-ATPase inhibitor. Each inhibitor effectively blocked endosomal acidification as evidenced by decrease in Lysotracker fluorescence in inhibitor-treated cells (Fig. S9E). Entry of VSVΔG-G, which requires low pH as a trigger for membrane fusion, was sensitive to all three inhibitors in both cell lines (Figs. S9A and C). By contrast, VSVΔG-PIV5 entry, which occurs by fusion at the plasma membrane, was insensitive to any of the inhibitors (Figs. S9B and D).

VSVΔG-BHLD entry into C10 cells was inhibited by all three endosomal acidification inhibitors (Fig. 7B) and thus appears to require endosomal acidification. VSVΔG-BHLD entry into CHO-HVEM cells was inhibited only by one out of three inhibitors, NH_4_Cl (Fig. 7D) and likely does not require endosomal acidification. Interestingly, HSV-1 entry into both C10 and CHO-HVEM cells was inhibited by NH_4_Cl and monensin but not by BFLA (Figs. 7A and C). We hypothesize that the discrepancy in the inhibitory effects among the three inhibitors could potentially be due to the distinct mechanisms by which BFLA, NH_4_Cl, and monensin raise endosomal pH.

**Fig. 7.**
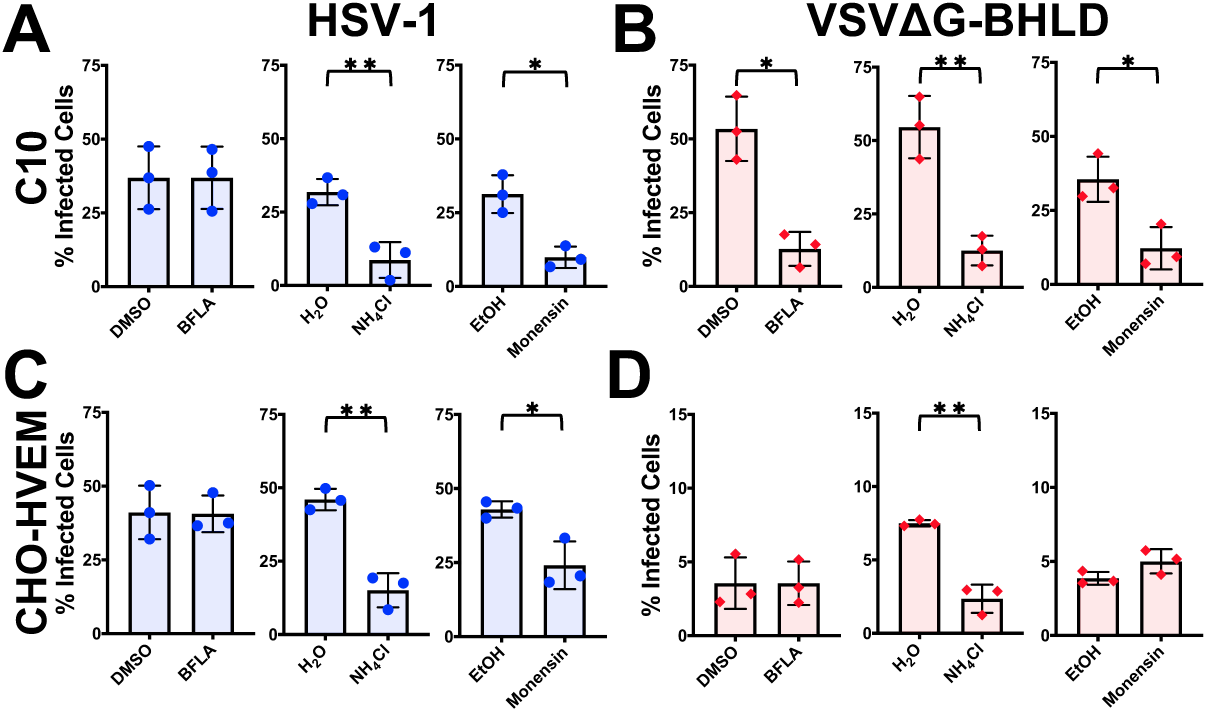
VSVΔG-BHLD entry requires endosomal acidification in a cell-dependent manner. C10 (A and B) and CHO-HVEM (C and D) cells were pretreated with inhibitors of endosomal acidification BFLA (100 nM), NH_4_Cl (50 mM), or monensin (15 μM) and infected with HSV-1 or VSVΔG-BHLD at MOI = 1. Infectivity was quantitated by flow cytometry at 6 hours post infection. Significance was calculated using a two-tailed Student’s T-test with Welch’s correction (p < 0.05 = *; p < 0.01 = **; p < 0.001 = ***).

### Glycoprotein C (gC) increases entry efficiency into CHO-HVEM and HaCaT cells

The much narrower tropism of VSVΔG-BHLD pseudotype relative to HSV-1 suggested that envelope proteins outside the essential four may contribute to HSV-1 tropism and entry efficiency. To test this hypothesis, we generated a VSV-pseudotype containing gC in addition to gB, gH, gL, and gD (VSVΔG-BHLD-gC) (Fig. 8A). To generate the VSVΔG-BHLD-gC pseudotype at sufficiently high titers suitable for entry experiments, the amount of gB plasmid transfected into HEK293T cells was increased. The corresponding VSVΔG-BHLD-pCAGGS control was generated similarly. For yet unclear reasons, VSVΔG-BHLD-pCAGGS entered CHO-nectin-1, CHO-HVEM, and HaCaT cells (Fig. 8B) more efficiently than the VSVΔG-BHLD pseudotype (Fig. 1A). Transfection of higher amounts of the gB plasmid could, in principle, lead to a higher expression levels of gB in the cells and, consequently, higher incorporation into the virions. More importantly, however, the VSVΔG-BHLD-gC pseudotype entered CHO-HVEM and HaCaT cells with a significantly higher efficiency than the VSVΔG-BHLD-pCAGGS pseudotype (Fig. 8B). These results suggest that gC can increase cell entry efficiency in a cell-specific manner.

**Fig. 8.**
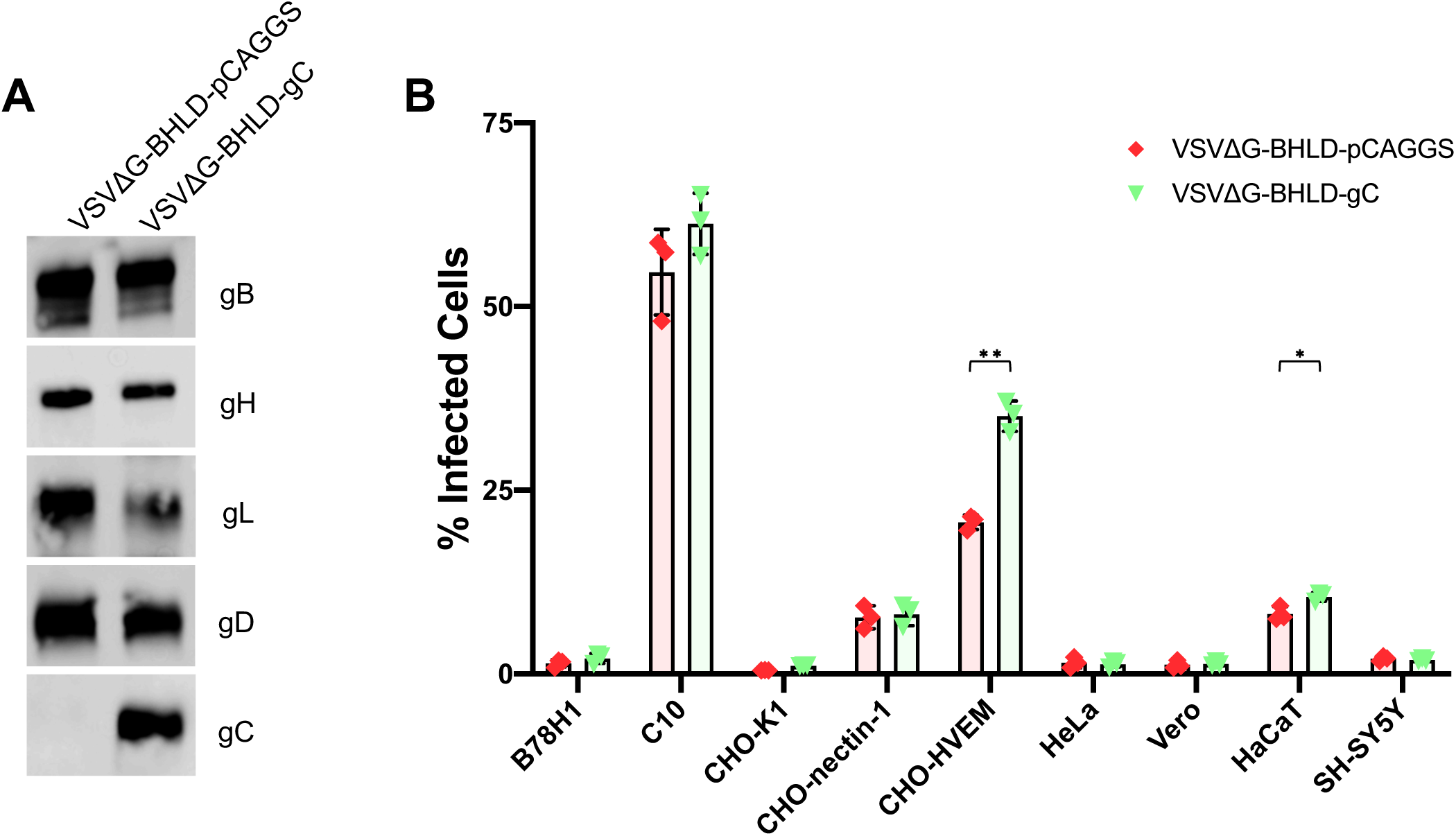
gC increases entry efficiency into CHO-HVEM and HaCaT cells. A) Incorporation of gC was verified by western blot of pelleted and washed virions. B) VSVΔG-BHLD-pCAGGS and VSVΔG-BHLD-C entry was assessed on nine cell lines, B78H1, C10, CHO-K1, CHO- nectin-1, CHO-HVEM, HeLa, Vero, HaCaT, and SH-SY5Y. Cells were infected at a MOI of 1. Entry was quantitated by flow cytometry at 6 hours post infection. Significance was calculated using a two-tailed Student’s T-test with Welch’s correction (p < 0.05 = *; p < 0.01 = **; p < 0.001 = ***).

## Discussion

Decades ago, glycoproteins gB, gH, gL, and gD were established as essential for HSV-1 entry (6–8). These four glycoproteins are also sufficient for cell-cell fusion when co-expressed in uninfected, receptor-bearing cells (11, 12). While these studies greatly increased our understanding of the HSV-1 entry and fusion mechanisms, it was unclear whether these four glycoproteins were sufficient to specify cellular tropism and the selection of entry routes, partly due to the presence of up to 12 other envelope proteins. To begin addressing this, we generated a VSV-based pseudotype containing HSV-1 gB, gH, gL, and gD. Being devoid of other HSV-1 proteins, the VSVΔG-BHLD pseudotype provides a bare-bones platform to identify contributions of the core set of four essential glycoproteins to HSV-1 cellular tropism and the selection of entry routes.

Previously, we showed that the VSVΔG-BHLD pseudotype efficiently entered C10 cells and that its entry recapitulated several important features of HSV-1 entry into susceptible cells: the requirement for gB, gH, gL, gD, and a gD receptor and sensitivity to anti-gB and anti-gH/gL neutralizing antibodies (39). Here, we expanded this study to six additional HSV-1-susceptible cell lines and made two key observations. First, we found that in addition to C10 cells, only CHO-HVEM cells supported appreciable VSVΔG-BHLD entry. Second, VSVΔG-BHLD and HSV-1 entered these two cell lines by distinct endocytic mechanisms as judged by the differences in sensitivity to various inhibitors (Fig. 9 and Table S1). These results imply that alone, gB, gH, gL, and gD permit entry of VSV pseudotypes only into a limited range of HSV-1-susceptible cell types and even then, do not specify native entry routes. On the basis of these results, we hypothesize that other HSV-1 envelope proteins may have underappreciated roles in defining HSV-1 tropism, entry route selection, or both. Although it may be too early to conclude that the incorporation of gC into the VSVΔG-BHLD pseudotype has changed its tropism, the increase in cell-specific entry efficiency of the VSVΔG-BHLD-gC pseudotype supports the use of the pseudotyping platform developed here for future gain-of-function studies. In these future studies, changes in tropism, entry routes, or both, may be uncovered.

**Fig. 9.**
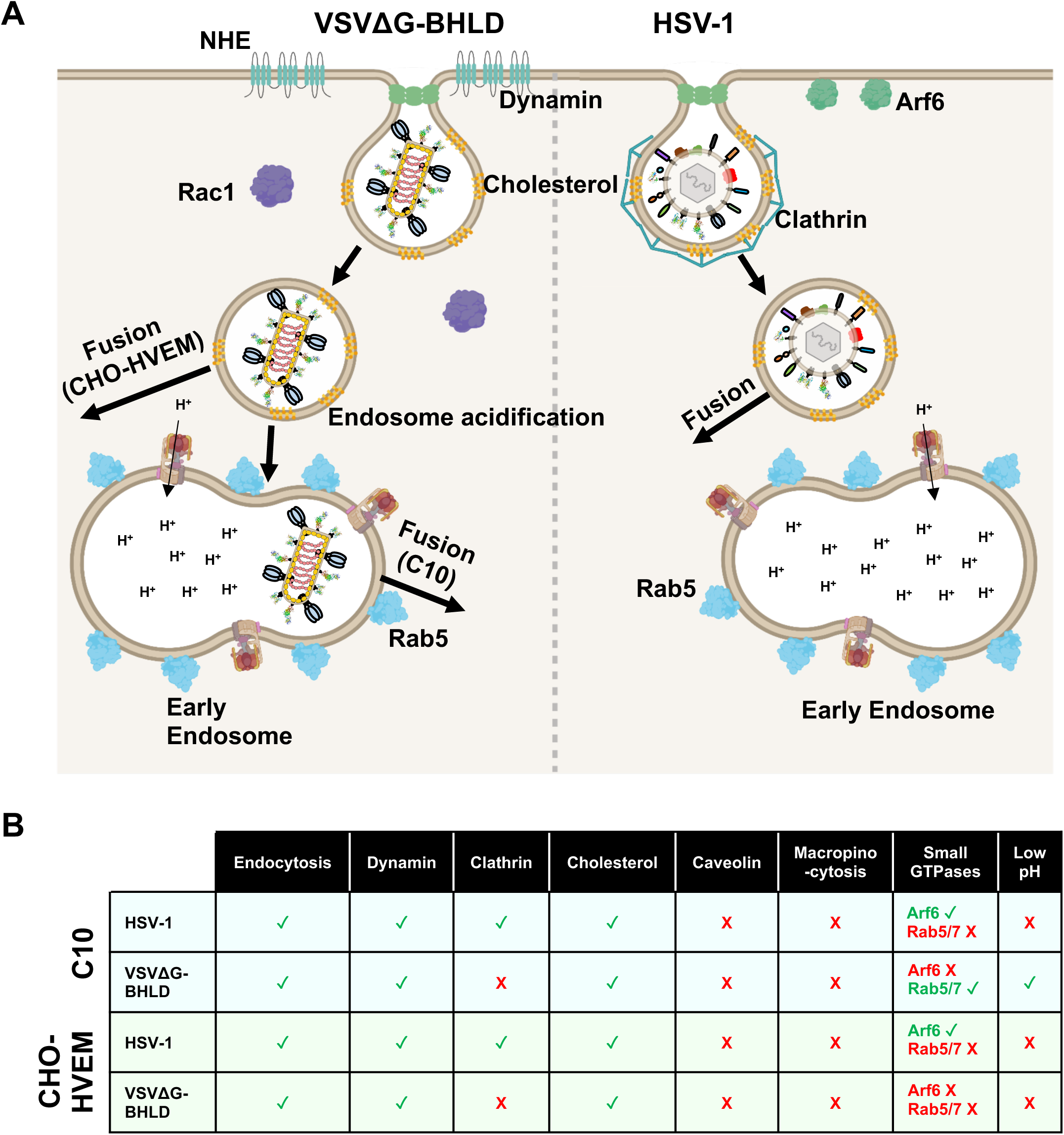
Entry model of VSVΔG-BHLD and HSV-1. A) Entry of VSVΔG-BHLD and HSV-1 entry into C10 and CHO-HVEM cells occurs by endocytosis and requires dynamin and cholesterol. VSVΔG-BHLD entry into C10 and CHO-HVEM cells additionally requires NHE and Rac1 activity whereas HSV-1 does not. VSVΔG-BHLD entry into C10 cells, but not CHO-HVEM cells also requires Rab5 and endosomal acidification. HSV-1 does not require either Rab5 or endosomal acidification for entry into either cell type. B) Table summarizing the cellular molecules important for HSV-1 and VSVΔG-BHLD entry into C10 and CHO-HVEM cells. Green check marks indicate that the virus requires that cellular component for entry. Red X- marks indicate that the virus does not require that cellular component for entry.

### VSVΔG-BHLD pseudotype has a narrower cellular tropism than HSV-1

HSV-1 can infect a wide range of receptor-bearing cell types from different species [reviewed in (25)]. However, the VSVΔG-BHLD pseudotype has a narrower tropism, efficiently entering only 2 out of 7 tested cell lines. Puzzlingly, while VSVΔG-BHLD entered two engineered rodent cell lines, it exhibited little to no entry into the four human and primate cell lines typically used in HSV-1 studies, even at an MOI of 10. The lack of VSVΔG-BHLD entry did not correlate with the HSV-1 entry route into these cells, namely, endocytosis for HeLa and HaCaT (5, 46) vs. plasma membrane for Vero and SH-SY5Y (5, 44). Additionally, there was no clear correlation with the receptor type (nectin-1 vs. HVEM) or with the cell type. While any of these factors – species, cell type, receptor, or route of entry – could potentially account for the decreased entry efficiency observed for VSVΔG-BHLD pseudotype, none stood out as major infectivity determinants.

We do not yet fully understand the reasons for the observed differences in tropism between the VSVΔG-BHLD pseudotype and HSV-1. At a first glance, the differences in virion structure – bullet-shaped vs. spherical – could be responsible for the phenotypic differences. However, the apparent differences in the entry of VSVΔG-BHLD, VSVΔG-G, and VSVΔG-PIV5 pseudotypes, all of which share the same VSV structure, suggest that virion structure is unlikely to be a major factor responsible for the observed differences in entry of VSVΔG-BHLD and HSV-1. The gB:gH:gL:gD ratios were also similar between VSVΔG-BHLD and HSV-1. The two viruses could, however, differ in lipid composition because VSV and HSV-1 acquire their envelopes from different sources. VSV buds at the plasma membrane (PM) (27) whereas HSV-1 buds at the trans-Golgi network (TGN) (42) or endosome-derived vesicles (43). However, according to recent lipidomics studies, the PM, the TGN, and the endosomes have similar lipid compositions (84). While we acknowledge that even small differences in lipid composition of the envelope could potentially contribute to differences in entry routes, this line of inquiry is beyond the scope of the present study. Moreover, no benchmarks are in place because HSV-1 lipid composition is unknown and cannot be altered on demand.

Importantly, VSV pseudotypes have been successfully used to study the entry of enveloped viruses regardless of the envelope origin. In addition to viruses that, like VSV, acquire their envelopes from the plasma membrane (Ebola virus, Lassa virus, Lujo virus) (31–34), viruses that derive their envelopes from the ER (Hepatitis C virus, Japanese encephalitis virus) (37, 38) or the Golgi (Hantavirus, Rift Valley fever virus) (35, 36) have also been studied. These observations suggest that VSV pseudotypes can provide important insights into viral entry mechanisms regardless of envelope origins.

This leaves differences in glycoprotein content as a potential reason for the differences in tropism. HSV-1 has up to 12 envelope proteins outside of the core set of four, which are absent from VSVΔG-BHLD. We hypothesize that efficient entry by HSV-1 into susceptible cells requires one or more of these other proteins. Indeed, some of them have already been shown to increase HSV-1 entry efficiency. For example, deletion of the N terminus of glycoprotein K (gK) promoted inefficient endocytic entry into Vero cells (85, 86), which normally support entry by fusion at the plasma membrane (44). The N terminus of gK may thus regulate the fusion of the viral envelope with the plasma membrane (85). Another HSV-1 glycoprotein, gC, aids viral attachment by binding heparan sulfate moieties of cell surface proteoglycans (87) and promotes efficient entry into cells that HSV-1 enters by an endocytic route (88). Thus, envelope proteins outside of the core set of four could, indeed, modulate HSV-1 tropism by tuning entry efficiency. In other words, the more efficiently HSV-1 enters a given cell type, the more likely that cell type is to be successfully infected.

This hypothesis could be tested by adding the “non-essential” envelope proteins one-by-one into the VSVΔG-BHLD pseudotype to test for their ability to restore entry into specific cell lines. For example, the incorporation of gK into the VSVΔG-BHLD pseudotype (VSVΔG-BHLD-gK) would be expected to increase the entry efficiency into cells that HSV-1 enters by fusion at the plasma membrane (Vero and SH-SY5Y), whereas gC (VSVΔG-BHLD-gC) could increase the entry efficiency into cells that HSV-1 enters by endocytosis (C10, CHO-nectin-1, CHO-HVEM, HeLa and HaCaT). Indeed, we found that incorporating gC into the VSVΔG-BHLD pseudotype increased entry efficiency into CHO-HVEM and HaCaT cells. These results are consistent with the reduced entry efficiency of an HSV-1 mutant lacking gC into these cell types (88). The mechanism underlying the gC-dependent gain-of-function phenotype of VSVΔG-BHLD-gC will be explored in future work.

### VSVΔG-BHLD pseudotype is internalized differently from HSV-1

If HSV-1 gB, gH, gL, and gD were sufficient to specify the native routes of HSV-1 entry, then we would have expected the VSVΔG-BHLD pseudotype to utilize the same entry routes into C10 and CHO-HVEM as HSV-1. However, while entry of both viruses occurred by endocytosis and required dynamin and cellular cholesterol but not caveolin-1 or actin polymerization, further investigation uncovered several notable differences in entry requirements (Fig. 9).

The first difference was that HSV-1 entry into C10 or CHO-HVEM cells was inhibited by the clathrin inhibitor Pitstop-2, suggesting that it occurred by CME, whereas VSVΔG-BHLD entry was not. HSV-1 entry inhibition by Pitstop-2 was unexpected because HSV-1 does not appear to utilize clathrin for entry into several cell lines, including HaCaT (89), CHO-nectin-1 (90), Vero, HeLaS3, and HeLaCNX cells (91). However, the role of clathrin in entry into C10 or CHO-HVEM cells had not been assessed prior to this study. Moreover, recent work has suggested that HSV-1 entry into a human oligodendrocytic cell line (HOG) depends on clathrin (92). Therefore, HSV-1 may utilize CME in a cell-specific manner.

Unlike HSV-1 entry, VSVΔG-BHLD entry into C10 or CHO-HVEM cells was not inhibited by Pitstop-2, which implicated CIE, rather than CME, as the entry route. Macropinocytosis is a common CIE used by several viruses, but VSVΔG-BHLD entry into both C10 and CHO-HVEM cells was insensitive to the inhibitor of actin polymerization, cytochalasin D, as was HSV-1. Given the essential role of actin polymerization in macropinocytosis (68, 71), these data suggest that macropinocytosis is not the primary entry mechanism for VSVΔG-BHLD pseudotype or HSV-1. Accordingly, VSVΔG-BHLD particles did not colocalize with a fluid phase uptake marker 70 kDa rhodamine-B-labeled dextran to an appreciable extent. The lack of actin involvement in HSV-1 entry was not entirely surprising because its requirement, as deemed by cytochalasin D treatment, varies from cell line to cell line. For example, cytochalasin D treatment blocked entry into CHO-nectin-1 cells (90) but not into primary keratinocytes or HaCaT cells (89).

Unexpectedly, VSVΔG-BHLD entry into both C10 and CHO-HVEM cells was sensitive to two other inhibitors of macropinocytosis, EIPA and NSC23766. If the VSVΔG-BHLD pseudotype does not enter cells by macropinocytosis, why is its entry sensitive to EIPA and NSC23766? One possibility is that the respective targets of these inhibitors, Na^+^/H^+^ exchangers and Rac1, could contribute to VSVΔG-BHLD entry independently of their roles in macropinocytosis. For example, EIPA inhibits the function of other cellular GTPases like Rac1 and Cdc42 (69) whereas NSC23766 could affect other downstream targets of Rac1 [reviewed in (93)]. Alternatively, EIPA and NSC23766 could inhibit VSVΔG-BHLD entry due to their documented pleotropic effects on the cell. EIPA treatment can lead to a gross reorganization of the endosomal network and changes in Na^+^ and H^+^ gradients in the cell (94, 95). Similarly, Rac1, the target of NSC23766, is involved in several cellular processes in addition to regulating the actin cytoskeleton (96).

### VSVΔG-BHLD pseudotype and HSV-1 differ in late-stage entry requirements

Many viruses that enter by endocytosis, for example, influenza A and VSV, rely on Rab-GTPase-dependent endosomal maturation and acidification (74). VSVΔG-BHLD entry into C10 cells required Rab5, an early endosome marker, and endosomal acidification. VSVΔG-BHLD entry into CHO-HVEM cells and HSV-1 entry into both C10 and CHO-HVEM cells did not require either Rab5, or Rab7, or endosomal acidification. Nevertheless, efficient entry of HSV-1 into both C10 and CHO-HVEM cells required Arf6, a small GTPase involved in endosomal trafficking, including CME and CIE (79). How Arf6 promotes HSV-1 entry is yet unclear considering its numerous downstream effectors, including lipid modifying enzymes, proteins involved in endosome trafficking, GTPase activating proteins (GAPs) and guanine exchange factors (GEFs) for other GTPases [reviewed in (79)]. In contrast, Arf6 was dispensable for VSVΔG-BHLD entry.

One notable difference between VSVΔG-BHLD and HSV-1 entry into C10 cells was that VSVΔG-BHLD entry required endosomal acidification. Previous work suggested that HSV-1 entry into C10 cells did not require endosomal acidification (45). Indeed, we confirmed that HSV-1 entry into C10 cells was insensitive to bafilomycin A1 (BFLA), a well-known inhibitor of endosomal acidification. The requirement for endosomal acidification for VSVΔG-BHLD entry into C10 cells was unexpected. The membrane fusion itself may not require low pH, in agreement with the observations that cell-cell fusion in the presence of gB, gH, gL, and gD occurs at neutral pH (13). However, concomitant with the endosomal acidification, there are significant changes in endosomal ion concentrations and lipid content (97), which could affect membrane fusion or the fusion pore expansion. Therefore, we hypothesize that during VSVΔG-BHLD entry, endosomal acidification promotes the establishment of endosomal conditions conducive to fusion and that in HSV-1, envelope proteins outside the essential four may functionally replace endosomal acidification.

Surprisingly, HSV-1 entry into both C10 and CHO-HVEM cells was inhibited by two other inhibitors of endosomal acidification, NH_4_Cl and monensin. Previous work showed that HSV-1 entry into CHO-HVEM cells was sensitive to inhibitors of endosomal acidification ammonium chloride (NH_4_Cl) and monensin (46, 98), indicating a requirement for endosomal acidification. While NH_4_Cl and monensin alkalinize the lumen of endosomes, the mechanisms by which they do so differ dramatically from BFLA. NH_4_Cl, when dissolved, exists in equilibrium as NH_3_ and NH_4_^+^. Upon entering acidic environment, e.g., an endosome, NH_3_ becomes protonated to NH_4_^+^, which leads to an increase in endosomal pH (97). Monensin is a carboxylic ionophore that utilizes an electroneutral exchange of monovalent cations for protons, effectively raising the endosomal pH (97). In parallel to these alkalinizing effects, NH_4_Cl and monensin can affect other cellular processes, e.g., vacuolization or organelle swelling. In contrast, BFLA functions by specifically blocking the function of the V_0_ domain of V-ATPases thereby blocking the movement of protons across the endosomal membrane (99). At the nM concentrations used, BFLA is very specific and potent in its action. Therefore, given that HSV-1 entry into either C10 or CHO-HVEM cells was not blocked by BFLA, we hypothesize that it does not require endosomal acidification. Sensitivity of HSV-1 entry to NH_4_Cl and monensin could, instead, be due to their ability to interfere with other cellular processes. Both compounds alter ion content of the endosomes and cause vacuolization (97). While the impact of endosome vacuolization on HSV-1 entry has not been investigated, a change in endosomal ion concentration could, potentially, reduce the ability of HSV-1 to fuse with the endosomal membrane. Indeed, binding of HSV-1 to the cell surface releases intracellular Ca^2+^ stores (100) and increases intracellular levels of Cl^-^ ions (101), both of which appear important for subsequent entry.

Collectively, we hypothesize that HSV-1, which does not require Rab5/7 or endosomal acidification, fuses with the endosomal membrane prior to maturation of the newly formed vesicle into an early endosome [pH ∼6.2 (102)] (Fig. 9). This latter scenario is consistent with the rapid nature of HSV-1 entry into both C10 and B78A10 cells (B78 murine melanoma cells expressing HVEM) cells (t_1/2_ = 8-10 minutes) (45). Alternatively, endosomal maturation status does not influence HSV-1 fusion with the membrane of the endocytic vesicle. VSVΔG-BHLD entry into CHO-HVEM cells, likewise, does not require Rab5/7 or endosomal acidification, implying that VSVΔG-BHLD may fuse with the endosomal membrane prior to delivery of the endocytic vesicle to an early endosome. By contrast, VSVΔG-BHLD entry into C10 cells requires both Rab5 and endosomal acidification, which suggests that VSVΔG-BHLD fuses with membranes of early endosomes. This would suggest that during HSV-1 entry into C10 cells, other envelope proteins may enable fusion prior to endosomal acidification.

As this and other studies show, HSV-1 entry is a complex phenomenon that requires at least four glycoproteins (gB, gH, gL, and gD) that operate in the presence of up to 12 additional envelope proteins, understudied with regard to entry. By establishing a platform where the functionality of the four essential HSV-1 entry glycoproteins could be evaluated in isolation, we demonstrated that they are insufficient to define HSV-1 tropism or specify native entry routes. We have expanded the use of this platform by demonstrating that incorporation of an additional envelope protein, gC, can lead to increased entry. Collectively, our work implicates other HSV-1 envelope proteins as underappreciated, yet potentially important contributors to HSV-1 tropism, entry route selection, and, ultimately, pathogenesis.

## Materials and Methods

### Cells

HEK293T (gift from John Coffin, Tufts University), Vero (ATCC® CCL-81™), HeLa (ATCC® CCL-2™), and HaCaT cells (gift from Jonathan Garlick, Tufts University) were grown in Dulbecco’s modified Eagle medium (DMEM; Lonza) containing high glucose, and sodium pyruvate, supplemented with L-glutamine (Caisson Labs), 10% heat inactivated fetal bovine serum (HI-FBS; Life Technologies) and 1X penicillin/streptomycin (pen/strep) solution (Corning). B78H1 cells (a gift from Gary Cohen, University of Pennsylvania) were grown in DMEM containing high glucose, sodium pyruvate, and L-glutamine supplemented with 5% FBS and pen/strep solution (1X). C10 cells (a gift from Gary Cohen, University of Pennsylvania), a clonal B78H1-derivative stably expressing human nectin-1, were grown in DMEM containing high glucose, sodium pyruvate, and L-glutamine supplemented with 5% FBS and pen/strep solution (1X) and maintained under selection for nectin-1 expression with 250 μg/ml of G418 (Selleck Chemical) as done previously (39). CHO-K1 cells were grown in Ham’s F12 medium containing 10% FBS and pen/strep solution (1X). CHO-HVEM cells, a derivative of CHO-K1 cells that stably express human HVEM, were grown in Ham’s F12 medium containing 10% FBS and penicillin-streptomycin solution (1X), 250 ug/ml G418 and 150 ug/ml of puromycin (AG Scientific). CHO-K1 and CHO-HVEM cells were a gift from Anthony Nicola (Washington State University). CHO-nectin-1 cells, a derivative of CHO-K1 cells that stably express human nectin-1 were grown in Ham’s F12 medium containing 10% FBS and penicillin-streptomycin solution (1X), 250 ug/ml G418 and 5 ug/ml of puromycin (AG Scientific). CHO-nectin-1 cells were a gift from Richard Longnecker (Northwestern University). SH-SY5Y cells were maintained in EMEM (Sigma-Aldrich) supplemented with 15% HI-FBS and 1X penicillin/streptomycin. SH-SY5Y cells were a kind gift from Stephen Moss (Tufts University).

### Plasmids

Plasmids pPEP98, pPEP99, pPEP100, and pPEP101 carry the full-length HSV-1 (strain KOS) genes for gB, gD, gH, and gL, respectively in a pCAGGS vector background. These were kindly gifted by P.G. Spear (Northwestern University). pCMV-VSV-G, which contains the full-length gene for the VSV glycoprotein, G, was a gift from Judith White (University of Virginia). Rab GTPase dominant negative constructs [mCherry-Rab5DN(S34N) and dsRed-Rab7DN] were purchased from Addgene (76, 77). For consistency, the dsRed in dsRed-Rab7DN was replaced with mCherry by amplifying mCherry with the following primers: 5’-AGCGCTACCGGTCGCCACCATGGTGAGCAAGGGCGAG-3’ (forward) and 5’- AATTCGAAGCTTGAGCTCGAGATCTGAGCTTGTACAGCTCGTCCATGCC-3’ (reverse). mCherry was then cloned in frame with Rab7DN using AgeI and HindIII cut sites that were engineered into the forward and reverse primers, respectively. As our HSV-1 reporter strain uses tdTomato, eGFP-RabDN constructs were engineered. The same primers were used to amplify eGFP from pEGFP-N2. The same cloning procedure was used to replace mCherry and dsRed with eGFP in the Rab5DN and Rab7DN constructs, respectively. Isolated clones were sequenced to verify mCherry and eGFP were in frame with the DN Rab genes. HSV-1 gC was amplified from HSV-1 (F strain) BAC DNA (GS6000) with the following primers: 5’- CGAGCTCGGCCACCATGGCCCCGGGGCGGG-3’ (forward) and 5’- GGGGTACCCCTCACGTAGAATCGAGACCGAGGAGAGGGTTAGGGATAGGCTTACCCCGCCGATGACGCTGCCG-3’ (reverse). The amplicon was digested with SacI and KpnI and cloned into the expression vector, pCAGGS. The C-terminus of gC was tagged with a V5 epitope tag for western blot purposes.

### Antibodies

Nectin-1 antibody [clone CK41 (103)] conjugated to phycoerythrin (PE) was purchased from BD Biosciences. PE-isotype antibody was also purchased from BD Biosciences. HVEM antibody (R140) was a gift from Gary Cohen (University of Pennsylvania). Caveolin-1 antibody (clone 4H312) was purchased from Santa Cruz Biotechnology. β-actin antibody conjugated to horse radish peroxidase(sc-47778 HRP) was purchased from Santa Cruz Biotechnology. Anti-V5 antibody (V8137) was purchased from Sigma-Aldrich.

### Chemical inhibitors

Monensin, methyl-β-cyclodextrin, cytochalasin D, Pitstop-2, and EIPA were purchased from Sigma. Dynasore and MiTMAB were purchased from Calbiochem. Bafilomycin A1 was purchased from ApexBio. Ammonium chloride was purchased from Fisher Scientific. Dyngo-4a was purchased from Abcam. NSC23766 was purchased from Santa Cruz Biotechnology.

### Viruses

Pseudotyped viral particles (VSVΔG-BHLD) were generated as described previously (39). Briefly, HEK293T cells (5.5 x 10^6^ cells/10 cm dish) were transfected with 2.5 μg each pPEP98, pPEP99, pPEP100, and pPEP101 using polyethyleneimine (PEI at 1 mg/ml) at a 3:1 weight ratio of PEI to DNA. VSVΔG-BHLD-pCAGGS and VSVΔG-BHLD-gC were generated by transfecting HEK293T cells with 10 μg pPEP98 and 2.5 μg each of pPEP99, pPEP100, pPEP101, and pCAGGS or pCAGGS-gC-V5 using GenJet Ver. II (SignaGen Laboratories). In all cases, 24 hours post transfection, cells were infected at an MOI = 3 with VSVΔG-G (VSVΔG pseudotyped with VSV G protein) and incubated at 30° C. Forty-eight hours post infection, supernatants were collected, cleared of cell debris (two spins at 1500 x g for 10 minutes each), and stored at −80° C. VSVΔG-BHLD, VSVΔG-BHLD-pCAGGS, and VSVΔG-BHLD-gC titers were determined on C10 cells.

HSV-1 (GS3217, F strain) was kindly provided by Gregory Smith (Northwestern University). GS3217 contains a tdTomato reporter gene with a nuclear localization signal under control of a CMV immediate early (IE) promoter (104). HSV-1 was propagated on Vero cells, and titers were determined by plaque assay on Vero cells as previously described (105). VSVΔG-G helper virus was generated by Michael Whitt (University of Tennessee) and kindly provided by Judith White (University of Virginia). New stocks were generated similarly to the VSVΔG-BHLD pseudotypes, replacing the HSV-1 glycoproteins with pCMV-VSV-G (10 μg per 10 cm dish). VSVΔG-G titers were determined on C10 cells. VSVΔG-PIV5 was generated and kindly provided by S.P.J. Whelan (Washington University). As VSVΔG-PIV5 contains the PIV5 HN and F proteins in the VSV genome, no complementation *in trans* was necessary. VSVΔG-PIV5 was grown on HEK293T cells and titers were determined on C10 cells. Entry of the VSV pseudotypes (VSVΔG-BHLD, VSVΔG-G, VSVΔG-PIV5, VSVΔG-BHLD-pCAGGS, and VSVΔG-BHLD-gC) was assessed by the expression of the GFP reporter driven by the promoter within the 3’ leader sequence of the VSV genome.

### Entry experiments

3×10^5^ B78H1, C10, CHO-K1, CHO-nectin-1, CHO-HVEM, HeLa, HaCaT, Vero, or SH-SY5Y cells were seeded in 35 mm dishes. Cells were infected with viruses at a MOI=1. Viruses were incubated with cells at 37° C for one hour. After one hour, viruses that had not entered were inactivated with a low pH wash (40 mM Na citrate, 10 mM KCl, 135 mM NaCl, pH 3.0). Complete growth media was added back to cells and infections were allowed to progress for six hours prior to analysis by flow cytometry. Entry experiments in the presence of inhibitors were performed similarly except that prior to infection, C10 and CHO-HVEM cells were pretreated with the indicated inhibitors for one hour prior to infection. All inhibitors, with the exception of sucrose and methyl-β-cyclodextrin, were present during the infection and the six hours post infection prior to analysis by flow cytometry by measuring tdTomato expression (HSV-1 GS3217) or EGFP expression (VSV pseudotypes) to allow sufficient time for viral entry and expression of the fluorescent reporters.

Viral entry experiments in the presence of fluorescently labeled Rab GTPase dominant negative constructs were performed and analyzed as follows: cells infected with HSV-1 (tdTomato) were transfected with pEGFP-N2 as an empty vector control or eGFP-tagged RabDN constructs whereas cells infected with VSV pseudotypes (eGFP) were transfected with pmCherry-C1 empty vector control or mCherry-tagged RabDN constructs. The data are represented as the percentage of infected and transfected cells out of the total number of transfected cells with either the empty vector control or the RabDN constructs.

Prior to flow cytometry analysis, cells were trypsinized, resuspended in media and pelleted at 450 x g for five minutes. Cells were washed with 1X PBS containing 1 mM EDTA (to prevent clumping). Cells were pelleted again at 450 x g for five minutes. Cells were then resuspended in 1X PBS with 1 mM EDTA and transferred to FACS tubes. Flow cytometry was performed on a BD LSR II or FACSCalibur instrument. tdTomato expression (HSV-1 GS3217) or EGFP expression (VSV pseudotypes) were measured as a proxy for viral entry. Data analysis was done using FlowJo software (v. 8.8.7).

### Analyses of receptor expression by flow cytometry

Nectin-1 was detected on the surface of cells by staining them with anti-nectin-1 antibody CK41 conjugated to PE (BD Biosciences). HVEM was detected on the surface of cells using the anti-HVEM polyclonal antibody, R140, a gift from Gary Cohen (University of Pennsylvania), and a FITC-conjugated anti-rabbit secondary antibody (ThermoFisher). Briefly, 1×10^6^ cells were plated into 10 cm dishes. The next day, cells were lifted from the dishes with 1X-PBS containing 5 mM EDTA. Cells were then pelleted, resuspended in 300 μl FACS buffer (1X-PBS, 2% FBS, 1 mM EDTA), and divided evenly between microfuge tubes for mock, PE-isotype, or PE-anti-nectin-1 treatment. Cells were incubated with 1 μg of antibody for 30 minutes on ice with agitation every 10 minutes. Similarly, cells were divided for mock, isotype (anti-gB R68 polyclonal antibody), or R140 anti-HVEM treatment. Cells were incubated with 5 μg of antibody. Mock, isotype, and R140-labeled cells were then incubated with a FITC-conjugated anti-rabbit secondary antibody (ThermoFisher). After 30 minutes, cells were pelleted and washed three times with FACS buffer, re-suspended, then immediately analyzed by flow cytometry (FACSCalibur).

### Virus purification and densitometry analysis

HSV-1 and VSVΔG-BHLD virions were purified and subjected to immunoblot for gB, gH, gL, and gD. Briefly, five T-175 flasks of Vero cells were infected with HSV-1 (MOI 0.01). HSV-1 was crudely purified as previously described (105). HSV-1 particles were then purified over a continuous 15-50% sucrose gradient (106). The purified band of HSV-1 was collected by puncture and aspiration. VSVΔG-BHLD virions were generated as previously mentioned (see *Viruses* section of Materials and Methods). VSVΔG-BHLD particles were then pelleted at 20,000 RPM. VSVΔG-BHLD virions were then resuspended and purified over a continuous 15-35% Optiprep gradient [protocol adapted from (107, 108)] and collected by puncture and aspiration. HSV-1 and VSVΔG-BHLD virions were pelleted at 20,0000 RPM. Western blots for gB, gH, gL, and gD were done using the rabbit polyclonal R68 antibody (gB), the rabbit polyclonal R137 antibody (gH), the mouse monoclonal antibody L1 (gL), and the rabbit polyclonal antibody R7 (gD). Secondary antibodies from LI-COR were used in order to perform densitometry analysis using the Image Studio Lite software (IRDye^®^ 680RD goat anti-rabbit and IRDye^®^ 800CW goat anti-mouse). Raw densitometry values for gH, gL, and gD blots were normalized to their respective raw densitometry values for gB and reported as fold-differences to gB.

### Confocal microscopy

1×10^5^ cells (C10 and CHO-HVEM) were seeded onto 12 mm glass coverslips (Chemglass) in 24 well plates. Prior to labeling cells with specific markers of different endocytosis pathways, cells were pretreated with inhibitors for one hour at 37 °C. Post pre-treatment, cells were chilled to 4 °C for 10-15 minutes and were subsequently incubated with specific endocytic markers: Transferrin-Alexa Fluor 488 (50 μg/ml, Thermo Fisher Scientific), 70-kD dextran-rhodamine B (1 mg/ml, Thermo Fisher Scientific), or Lysotracker (1 μM, Thermo Fisher Scientific). Cells were incubated with endocytic markers for 10 minutes at 4° C. After the 10-minute incubation, C10 cells were shifted to 37° C for 10 minutes and CHO-HVEM cells were shifted to 37° C for 30 minutes (Transferrin-Alexa Fluor 488 and Lysotracker) or 40 minutes (70 kDa dextran) (109). After incubation at 37° C, cells were washed 3 times with 1X-PBS and fixed in 4% paraformaldehyde for 15 minutes at room temperature. Cells were washed three times with 1X-PBS and incubated with 2.5 μg/ml of DAPI (ThermoFisher Scientific) diluted in 1X-PBS for 15 minutes at room temperature. Cells were washed again three times with 1X-PBS and mounted onto Prolong Gold Antifade (Life Technologies) on glass slides (Thermo Fisher Scientific). Coverslips were sealed with clear nail polish and analyzed by confocal microscopy using a Leica SPE microscope. Images were analyzed in Fiji (110). For 70-kDa dextran and VSVΔG-BHLD co-localization experiments, C10 and CHO-HVEM cells were incubated with 1 mg/ml 70 kDa dextran and VSVΔG-BHLD (MOI = 1) for 1 hour at 4° C to allow for virion attachment. Cells were then shifted to 37° C for 20 minutes. After 20 minutes, cells were prepared for confocal microscopy by fixing with 4% paraformaldehyde, permeabilized and blocked with 1X PBS containing 5% normal goat serum and 0.3% Triton X-100. Cells were then incubated with anti-gB antibody (R68) overnight at 4° C. The next day, cells were incubated with a secondary antibody labeled with FITC for 1 hour at room temperature. Slides were then prepared as described above.

### siRNA-mediated knockdown

Mouse caveolin-1 siRNA, and a control siRNA were purchased from Santa Cruz Biotechnologies. 50 pmol of siRNA (0.625 μg) (cav-1 or scramble [scr]) were diluted into 100 μl of Optimem. In another tube, 3.125 μl of PEI (1 mg/ml) was diluted into 100 μl of Optimem. The diluted PEI was mixed with the diluted siRNA to a final ratio of 5:1 (w:w) of PEI to siRNA and incubated at room temperature for 30 minutes. The complex was added dropwise to CHO-HVEM cells plated in 35 mm dishes (3 x 10^5^ cells/dish). Cells were incubated at 37° C for 48 hours before infection and subsequent flow cytometry analysis.

## Acknowledgments

We thank Stephen Kwok and Allen Parmelee at the Tufts Laser Cytometry Core facility for their help with FACS experiments; Gary Cohen (University of Pennsylvania) for the gifts of antibodies and cell lines; Sean Whelan (Washington University) for the gift of VSVΔG-PIV5 pseudotype; Anthony Nicola (Washington State University) for the gift of CHO-K1 and CHO-HVEM cells; Richard Longnecker (Northwestern University) for the gift of CHO-nectin-1 cells; John Coffin (Tufts University) for the gift of HEK293T cells; Stephen Moss (Tufts University) for the gift of SH-SY5Y cells; Jonathan Garlick (Tufts University) for the gift of HaCaT cells; Michael Forgac (Tufts University) for the gift of Lysotracker reagent; Gregory Smith (Northwestern University) for the gift of HSV-1 GS3217 strain, and Michael Whitt (University of Tennessee) and Judith White (University of Virginia) for the gift of the VSVΔG-GFP pseudotyping platform. FACS experiments were performed at the Tufts Laser Cytometry Core facility.

## Data sharing plan

All data is included in the manuscript and supporting information.

## Supplemental Figure Legends

**Fig. S1. Infecting cells with VSVΔG-BHLD at a higher MOI does not increase entry to an appreciable extent.** A)Receptor null (B78H1 and CHO-K1) and receptor bearing cells (C10, CHO-HVEM, HeLa, Vero, HaCaT, and SH-SY5Y) were infected at MOI =1 (red) or MOI = 10 (purple). Entry efficiency was assessed by flow cytometry at 6 hours post infection. B and C) Receptor null (B78H1 and CHO-K1) and receptor bearing cells (C10, CHO-HVEM, HeLa, Vero, HaCaT, and SH-SY5Y) were infected at MOI =1 with either VSVΔG-G (B) or VSVΔG-PIV5 (C). Entry was assessed by flow cytometry at 6 hours post infection.

**Fig. S2. VSVΔG-G and VSVΔG-PIV5 entry in the presence of hypertonic sucrose.** C10 (A, B) and CHO-HVEM (C, D) cells were pretreated with a hypertonic solution of sucrose (0.3 M) and infected with VSVΔG-G or VSVΔG-PIV5 at MOI = 1. Infectivity was quantitated by flow cytometry at 6 hours post infection. Significance was calculated using a two-tailed Student’s T-test with Welch’s correction (p < 0.05 = *; p < 0.01 = **; p < 0.001 = ***). E) C10 and CHO-HVEM cells were pretreated with 0.3 M sucrose and incubated with 50 ug/ml of AF488-labeled transferrin (Tf). Cells were fixed, counterstained with DAPI, and imaged by confocal microscopy. Scale bar = 25 µm.

**Fig. S3 VSVΔG-G and VSVΔG-PIV5 differ in their dependence on dynamin and clathrin for entry.** C10 (A and C) and CHO-HVEM (B and D) cells were pretreated with dynamin inhibitors Dynasore (80 μM), Dyngo-4a (25 μM), MiTMAB (5 μM), or the CME inhibitor Pitstop-2 (30 μM) and infected with VSVΔG-G or VSVΔG-PIV5 at a MOI of 1. Infectivity was quantitated by flow cytometry at 6 hours post infection. CHO-HVEM cells treated with Dyngo-4a or MiTMAB used the same DMSO control as indicated by the same bar graph appearing twice each in panels C and D. Significance was calculated using a two-tailed Student’s T-test with Welch’s correction (p < 0.05 = *; p < 0.01 = **; p < 0.001 = ***). E and F) C10 and CHO-HVEM cells were pretreated with dynamin inhibitors Dynasore, Dyngo-4a, MiTMAB, or Pitstop-2 at the same concentrations as in panels A-D and then incubated with 50 μg/ml of AF488-labeled transferrin. Cells were fixed, counterstained with DAPI, and imaged by confocal microscopy. Scale bar = 25 μm.

**Fig. S4. VSVΔG-G entry does not require cholesterol whereas VSVΔG-PIV5 entry requires cholesterol in a cell-type-dependent manner.** C10 (A and B) and CHO-HVEM (C and D) cells were pretreated with a cholesterol-removal drug methyl-β-cyclodextran, MβCD (5 mM) and infected with VSVΔG-G or VSVΔG-PIV5 at a MOI of 1. Infectivity was quantitated by flow cytometry at 6 hours post infection. E) C10 and CHO-HVEM cells were treated with either a solvent control (H_2_O/EtOH) or methyl-β-cyclodextrin (MβCD), then incubated with cholera toxin subunit B labelled with Alexa Fluor 488. Confocal microscopy was performed on the solvent control and methyl-β-cyclodextrin treated cells. Cells were fixed, counterstained with DAPI, and imaged by confocal microscopy. Scale bar = 25 μm. (F) CHO-HVEM cells were transfected with a caveolin-1 siRNA (cav-1) or a scrambled control siRNA (scr) (both 50 pm) and infected with VSVΔG-G or VSVΔG-PIV5 at a MOI of 1. Infectivity was quantitated by flow cytometry at 6 hours post infection. Significance was calculated using a two-tailed Student’s T-test with Welch’s correction (p < 0.05 = *; p < 0.01 = **; p < 0.001 = ***).

**Fig. S5. VSVΔG-BHLD does not co-localize with the fluid-phase marker 70 kDa dextran in C10 cells.** C10 cells were incubated with 1 mg/ml of rhodamine-B labelled 70 kDa dextran and VSVΔG-BHLD (MOI = 1) for one hour at 4° C. Cells were then shifted to 37° C for 20 minutes. Cells were fixed, counterstained with DAPI, and imaged by confocal microscopy. gB was detected by immunofluorescence using the rabbit pAb R68 and anti-rabbit IgG conjugated to FITC. Green = gB (marker for VSVΔG-BHLD particles); Red = 70 kDa dextran. Scale bar = 25 μm.

**Fig. S6. VSVΔG-BHLD, in large, does not co-localized with the fluid-phase marker, 70 kDa dextran, in CHO-HVEM cells.** CHO-HVEM cells were incubated with 1 mg/ml of rhodamine-B labelled 70 kDa dextran and VSVΔG-BHLD (MOI = 1) for one hour at 4°C. Cells were then shifted to 37° C for 20 minutes. Cells were fixed, counterstained with DAPI, and imaged by confocal microscopy. gB was detected by immunofluorescence using the rabbit pAb R68 and anti-rabbit IgG conjugated to FITC. Green = gB (marker for VSVΔG-BHLD particles); Red = 70 kDa dextran. Scale bar = 25 μm.

**Fig. S7. VSVΔG-G and VSVΔG-PIV5 entry does not require macropinocytosis.** C10 (A and B) and CHO-HVEM (C and D) cells were pretreated with macropinocytosis inhibitors cytochalasin D (2 μM), EIPA (25 μM), or NSC23766 (200 μM) and infected with VSVΔG-G or VSVΔG-PIV5 at a MOI of 1. Infectivity was quantitated by flow cytometry at 6 hours post infection. Significance was calculated using a two-tailed Student’s T-test with Welch’s correction (p < 0.05 = *; p < 0.01 = **; p < 0.001 = ***). E) C10 and CHO-HVEM cells were pretreated with macropinocytosis inhibitors cytochalasin D, EIPA, or NSC23766 at the same concentrations as in panels A-D and then incubated with 1.0 mg/ml of Rhodamine-B-labeled 70-kDa dextran (Dex). Cells were fixed, counterstained with DAPI, and imaged by confocal microscopy. Scale bar = 25 μm.

**Fig. S8: Roles of Rab5, Rab7, and Arf6 in VSVΔG-G and VSVΔG-PIV5 entry.** The roles of the small GTPases Rab5, Rab7 (A, C, E, G), and Arf6 (B, D, F, H) were assessed for VSVΔG-G (A, B, E, F) and VSVΔG-PIV5 (C, D, G, H) entry into C10 (A, B, C, D) and CHO-HVEM (E, F, G, H) cells. C10 (A and C) and CHO-HVEM (E and G) cells were transfected with either an empty vector control (eGFP or mCherry), eGFP or mCherry-tagged Rab5 dominant negative (DN), or eGFP or mCherry-tagged Rab7DN. Cells were infected at an MOI = 1 with either VSVΔG-G or VSVΔG-PIV5. Entry was assessed by flow cytometry at 6 hpi. The percent of infected cells was determined by dividing the number of virus(+)/eGFP/mCherry(+) cells by the total number of eGFP/mCherry(+) cells. C10 (B and D) and CHO-HVEM cells (F and H) were treated with the Arf6 inhibitor NAV-2729 (25 μM) and infected with either VSVΔG-G or VSVΔG-PIV5 at an MOI = 1. Significance was calculated using a two-tailed Student’s T-test with Welch’s correction (ns = not significant; p < 0.05 = *; p < 0.01 = **; p < 0.001 = ***).

**Fig. S9. VSVΔG-G but not VSVΔG-PIV5 entry requires endosomal acidification.** C10 (A and B) and CHO-HVEM (C and D) cells were pretreated with inhibitors of endosomal acidification BFLA (100 nM), NH_4_Cl (50 mM), or monensin (15 μM) and infected with VSVΔG-G or VSVΔG-PIV5 at MOI = 1. Infectivity was quantitated by flow cytometry at 6 hours post infection. Significance was calculated using a two-tailed Student’s T-test with Welch’s correction (p < 0.05 = *; p < 0.01 = **; p < 0.001 = ***). E) C10 and CHO-HVEM cells were pretreated with inhibitors of endosomal acidification at the same concentrations as in panels A-D (BFLA, NH_4_Cl, or monensin) and then incubated with Lysotracker (1 μM). Cells were fixed, counterstained with DAPI, and imaged by confocal microscopy. Scale bar = 25 μm.

**Table S1. Sensitivity of HSV-1, VSVΔG-BHLD, VSVΔG-G, and VSVΔG-PIV5 to specific inhibitors.** Green check marks indicate that virus entry is sensitive to that particular inhibitor. Red X marks indicate that the virus is not sensitive to that particular inhibitor.

## References

1. Steiner I, Benninger F. 2013. Update on herpes virus infections of the nervous system. Curr Neurol Neurosci Rep 13:414.

2. Bradley H, Markowitz LE, Gibson T, McQuillan GM. 2014. Seroprevalence of herpes simplex virus types 1 and 2--United States, 1999-2010. J Infect Dis 209:325–33.

3. Agelidis AM, Shukla D. 2015. Cell entry mechanisms of HSV: what we have learned in recent years. Future Virol 10:1145–1154.

4. Miranda-Saksena M, Denes CE, Diefenbach RJ, Cunningham AL. 2018. Infection and Transport of Herpes Simplex Virus Type 1 in Neurons: Role of the Cytoskeleton. Viruses 10.

5. Nicola AV, Hou J, Major EO, Straus SE. 2005. Herpes simplex virus type 1 enters human epidermal keratinocytes, but not neurons, via a pH-dependent endocytic pathway. J Virol 79:7609–16.

6. Cai WH, Gu B, Person S. 1988. Role of glycoprotein B of herpes simplex virus type 1 in viral entry and cell fusion. J Virol 62:2596–604.

7. Roop C, Hutchinson L, Johnson DC. 1993. A mutant herpes simplex virus type 1 unable to express glycoprotein L cannot enter cells, and its particles lack glycoprotein H. J Virol 67:2285–97.

8. Ligas MW, Johnson DC. 1988. A herpes simplex virus mutant in which glycoprotein D sequences are replaced by beta-galactosidase sequences binds to but is unable to penetrate into cells. J Virol 62:1486–94.

9. Heldwein EE, Krummenacher C. 2008. Entry of herpesviruses into mammalian cells. Cell Mol Life Sci 65:1653–68.

10. Connolly SA, Jackson JO, Jardetzky TS, Longnecker R. 2011. Fusing structure and function: a structural view of the herpesvirus entry machinery. Nat Rev Microbiol 9:369–81.

11. Muggeridge MI. 2000. Characterization of cell-cell fusion mediated by herpes simplex virus 2 glycoproteins gB, gD, gH and gL in transfected cells. J Gen Virol 81:2017–27.

12. Turner A, Bruun B, Minson T, Browne H. 1998. Glycoproteins gB, gD, and gHgL of herpes simplex virus type 1 are necessary and sufficient to mediate membrane fusion in a Cos cell transfection system. J Virol 72:873–5.

13. Atanasiu D, Saw WT, Cohen GH, Eisenberg RJ. 2010. Cascade of events governing cell-cell fusion induced by herpes simplex virus glycoproteins gD, gH/gL, and gB. J Virol 84:12292–9.

14. Eisenberg RJ, Atanasiu D, Cairns TM, Gallagher JR, Krummenacher C, Cohen GH. 2012. Herpes virus fusion and entry: a story with many characters. Viruses 4:800–32.

15. Spear PG, Eisenberg RJ, Cohen GH. 2000. Three classes of cell surface receptors for alphaherpesvirus entry. Virology 275:1–8.

16. Krummenacher C, Supekar VM, Whitbeck JC, Lazear E, Connolly SA, Eisenberg RJ, Cohen GH, Wiley DC, Carfi A. 2005. Structure of unliganded HSV gD reveals a mechanism for receptor-mediated activation of virus entry. EMBO J 24:4144–53.

17. Lazear E, Carfi A, Whitbeck JC, Cairns TM, Krummenacher C, Cohen GH, Eisenberg RJ. 2008. Engineered disulfide bonds in herpes simplex virus type 1 gD separate receptor binding from fusion initiation and viral entry. J Virol 82:700–9.

18. Cairns TM, Ditto NT, Atanasiu D, Lou H, Brooks BD, Saw WT, Eisenberg RJ, Cohen GH. 2019. Surface Plasmon Resonance Reveals Direct Binding of Herpes Simplex Virus Glycoproteins gH/gL to gD and Locates a gH/gL Binding Site on gD. J Virol 93.

19. Atanasiu D, Cairns TM, Whitbeck JC, Saw WT, Rao S, Eisenberg RJ, Cohen GH. 2013. Regulation of Herpes Simplex Virus gB-Induced Cell-Cell Fusion by Mutant Forms of gH/gL in the Absence of gD and Cellular Receptors. Mbio 4.

20. Gianni T, Amasio M, Campadelli-Fiume G. 2009. Herpes simplex virus gD forms distinct complexes with fusion executors gB and gH/gL in part through the C-terminal profusion domain. J Biol Chem 284:17370–82.

21. Atanasiu D, Whitbeck JC, de Leon MP, Lou H, Hannah BP, Cohen GH, Eisenberg RJ. 2010. Bimolecular complementation defines functional regions of Herpes simplex virus gB that are involved with gH/gL as a necessary step leading to cell fusion. J Virol 84:3825–34.

22. Chowdary TK, Cairns TM, Atanasiu D, Cohen GH, Eisenberg RJ, Heldwein EE. 2010. Crystal structure of the conserved herpesvirus fusion regulator complex gH-gL. Nat Struct Mol Biol 17:882–8.

23. Cooper RS, Heldwein EE. 2015. Herpesvirus gB: A Finely Tuned Fusion Machine. Viruses 7:6552–69.

24. Loret S, Guay G, Lippe R. 2008. Comprehensive characterization of extracellular herpes simplex virus type 1 virions. J Virol 82:8605–18.

25. Karasneh GA, Shukla D. 2011. Herpes simplex virus infects most cell types in vitro: clues to its success. Virol J 8:481.

26. Hilterbrand AT, Heldwein EE. 2019. Go go gadget glycoprotein!: HSV-1 draws on its sizeable glycoprotein tool kit to customize its diverse entry routes. PLoS Pathog 15:e1007660.

27. Whitt MA. 2010. Generation of VSV pseudotypes using recombinant DeltaG-VSV for studies on virus entry, identification of entry inhibitors, and immune responses to vaccines. J Virol Methods 169:365–74.

28. Fukushi S, Mizutani T, Saijo M, Matsuyama S, Miyajima N, Taguchi F, Itamura S, Kurane I, Morikawa S. 2005. Vesicular stomatitis virus pseudotyped with severe acute respiratory syndrome coronavirus spike protein. J Gen Virol 86:2269–2274.

29. Fukushi S, Watanabe R, Taguchi F. 2008. Pseudotyped vesicular stomatitis virus for analysis of virus entry mediated by SARS coronavirus spike proteins. Methods Mol Biol 454:331–8.

30. Nie J, Li Q, Wu J, Zhao C, Hao H, Liu H, Zhang L, Nie L, Qin H, Wang M, Lu Q, Li X, Sun Q, Liu J, Fan C, Huang W, Xu M, Wang Y. 2020. Establishment and validation of a pseudovirus neutralization assay for SARS-CoV-2. Emerg Microbes Infect 9:680–686.

31. Carette JE, Raaben M, Wong AC, Herbert AS, Obernosterer G, Mulherkar N, Kuehne AI, Kranzusch PJ, Griffin AM, Ruthel G, Dal Cin P, Dye JM, Whelan SP, Chandran K, Brummelkamp TR. 2011. Ebola virus entry requires the cholesterol transporter Niemann-Pick C1. Nature 477:340–3.

32. Jae LT, Raaben M, Riemersma M, van Beusekom E, Blomen VA, Velds A, Kerkhoven RM, Carette JE, Topaloglu H, Meinecke P, Wessels MW, Lefeber DJ, Whelan SP, van Bokhoven H, Brummelkamp TR. 2013. Deciphering the glycosylome of dystroglycanopathies using haploid screens for lassa virus entry. Science 340:479–83.

33. Jae LT, Raaben M, Herbert AS, Kuehne AI, Wirchnianski AS, Soh TK, Stubbs SH, Janssen H, Damme M, Saftig P, Whelan SP, Dye JM, Brummelkamp TR. 2014. Virus entry. Lassa virus entry requires a trigger-induced receptor switch. Science 344:1506–10.

34. Raaben M, Jae LT, Herbert AS, Kuehne AI, Stubbs SH, Chou YY, Blomen VA, Kirchhausen T, Dye JM, Brummelkamp TR, Whelan SP. 2017. NRP2 and CD63 Are Host Factors for Lujo Virus Cell Entry. Cell Host Microbe 22:688–696 e5.

35. Kleinfelter LM, Jangra RK, Jae LT, Herbert AS, Mittler E, Stiles KM, Wirchnianski AS, Kielian M, Brummelkamp TR, Dye JM, Chandran K. 2015. Haploid Genetic Screen Reveals a Profound and Direct Dependence on Cholesterol for Hantavirus Membrane Fusion. mBio 6:e00801.

36. Riblett AM, Blomen VA, Jae LT, Altamura LA, Doms RW, Brummelkamp TR, Wojcechowskyj JA. 2016. A Haploid Genetic Screen Identifies Heparan Sulfate Proteoglycans Supporting Rift Valley Fever Virus Infection. J Virol 90:1414–23.

37. Tani H, Komoda Y, Matsuo E, Suzuki K, Hamamoto I, Yamashita T, Moriishi K, Fujiyama K, Kanto T, Hayashi N, Owsianka A, Patel AH, Whitt MA, Matsuura Y. 2007. Replication-competent recombinant vesicular stomatitis virus encoding hepatitis C virus envelope proteins. J Virol 81:8601–12.

38. Tani H, Morikawa S, Matsuura Y. 2011. Development and Applications of VSV Vectors Based on Cell Tropism. Front Microbiol 2:272.

39. Rogalin HB, Heldwein EE. 2016. Characterization of Vesicular Stomatitis Virus Pseudotypes Bearing Essential Entry Glycoproteins gB, gD, gH, and gL of Herpes Simplex Virus 1. J Virol 90:10321–10328.

40. Robinson LR, Whelan SP. 2016. Infectious Entry Pathway Mediated by the Human Endogenous Retrovirus K Envelope Protein. J Virol 90:3640–9.

41. Krummenacher C, Baribaud F, Ponce de Leon M, Baribaud I, Whitbeck JC, Xu R, Cohen GH, Eisenberg RJ. 2004. Comparative usage of herpesvirus entry mediator A and nectin-1 by laboratory strains and clinical isolates of herpes simplex virus. Virology 322:286–99.

42. Henaff D, Radtke K, Lippe R. 2012. Herpesviruses exploit several host compartments for envelopment. Traffic 13:1443–9.

43. Hollinshead M, Johns HL, Sayers CL, Gonzalez-Lopez C, Smith GL, Elliott G. 2012. Endocytic tubules regulated by Rab GTPases 5 and 11 are used for envelopment of herpes simplex virus. EMBO J 31:4204–20.

44. Koyama AH, Uchida T. 1987. The mode of entry of herpes simplex virus type 1 into Vero cells. Microbiol Immunol 31:123–30.

45. Milne RS, Nicola AV, Whitbeck JC, Eisenberg RJ, Cohen GH. 2005. Glycoprotein D receptor-dependent, low-pH-independent endocytic entry of herpes simplex virus type 1. J Virol 79:6655–63.

46. Nicola AV, McEvoy AM, Straus SE. 2003. Roles for endocytosis and low pH in herpes simplex virus entry into HeLa and Chinese hamster ovary cells. J Virol 77:5324–32.

47. Dutta D, Donaldson JG. 2012. Search for inhibitors of endocytosis: Intended specificity and unintended consequences. Cell Logist 2:203–208.

48. Carpentier JL, Sawano F, Geiger D, Gorden P, Perrelet A, Orci L. 1989. Potassium depletion and hypertonic medium reduce “non-coated” and clathrin-coated pit formation, as well as endocytosis through these two gates. J Cell Physiol 138:519–26.

49. Bose S, Zokarkar A, Welch BD, Leser GP, Jardetzky TS, Lamb RA. 2012. Fusion activation by a headless parainfluenza virus 5 hemagglutinin-neuraminidase stalk suggests a modular mechanism for triggering. Proc Natl Acad Sci U S A 109:E2625–34.

50. Matlin KS, Reggio H, Helenius A, Simons K. 1982. Pathway of vesicular stomatitis virus entry leading to infection. J Mol Biol 156:609–31.

51. Superti F, Seganti L, Ruggeri FM, Tinari A, Donelli G, Orsi N. 1987. Entry pathway of vesicular stomatitis virus into different host cells. J Gen Virol 68 (Pt 2):387–99.

52. Johannsdottir HK, Mancini R, Kartenbeck J, Amato L, Helenius A. 2009. Host cell factors and functions involved in vesicular stomatitis virus entry. J Virol 83:440–53.

53. Marsh M, Helenius A. 2006. Virus entry: open sesame. Cell 124:729–40.

54. Mercer J, Schelhaas M, Helenius A. 2010. Virus entry by endocytosis. Annu Rev Biochem 79:803–33.

55. Kirchhausen T. 2000. Clathrin. Annu Rev Biochem 69:699–727.

56. Singh M, Jadhav HR, Bhatt T. 2017. Dynamin Functions and Ligands: Classical Mechanisms Behind. Mol Pharmacol 91:123–134.

57. Quan A, McGeachie AB, Keating DJ, van Dam EM, Rusak J, Chau N, Malladi CS, Chen C, McCluskey A, Cousin MA, Robinson PJ. 2007. Myristyl trimethyl ammonium bromide and octadecyl trimethyl ammonium bromide are surface-active small molecule dynamin inhibitors that block endocytosis mediated by dynamin I or dynamin II. Mol Pharmacol 72:1425–39.

58. Cureton DK, Massol RH, Saffarian S, Kirchhausen TL, Whelan SP. 2009. Vesicular stomatitis virus enters cells through vesicles incompletely coated with clathrin that depend upon actin for internalization. PLoS Pathog 5:e1000394.

59. Mayor S, Parton RG, Donaldson JG. 2014. Clathrin-independent pathways of endocytosis. Cold Spring Harb Perspect Biol 6.

60. Pelkmans L, Kartenbeck J, Helenius A. 2001. Caveolar endocytosis of simian virus 40 reveals a new two-step vesicular-transport pathway to the ER. Nat Cell Biol 3:473–83.

61. Xu Q, Cao M, Song H, Chen S, Qian X, Zhao P, Ren H, Tang H, Wang Y, Wei Y, Zhu Y, Qi Z. 2016. Caveolin-1-mediated Japanese encephalitis virus entry requires a two-step regulation of actin reorganization. Future Microbiol 11:1227–1248.

62. Williams TM, Lisanti MP. 2004. The caveolin proteins. Genome Biol 5:214.

63. de Almeida CJG. 2017. Caveolin-1 and Caveolin-2 Can Be Antagonistic Partners in Inflammation and Beyond. Front Immunol 8:1530.

64. Bender FC, Whitbeck JC, Ponce de Leon M, Lou H, Eisenberg RJ, Cohen GH. 2003. Specific association of glycoprotein B with lipid rafts during herpes simplex virus entry. J Virol 77:9542–52.

65. Chinnapen DJ, Chinnapen H, Saslowsky D, Lencer WI. 2007. Rafting with cholera toxin: endocytosis and trafficking from plasma membrane to ER. FEMS Microbiol Lett 266:129–37.

66. Danthi P, Chow M. 2004. Cholesterol removal by methyl-beta-cyclodextrin inhibits poliovirus entry. J Virol 78:33–41.

67. Dou X, Li Y, Han J, Zarlenga DS, Zhu W, Ren X, Dong N, Li X, Li G. 2018. Cholesterol of lipid rafts is a key determinant for entry and post-entry control of porcine rotavirus infection. BMC Vet Res 14:45.

68. Mercer J, Helenius A. 2009. Virus entry by macropinocytosis. Nat Cell Biol 11:510–20.

69. Koivusalo M, Welch C, Hayashi H, Scott CC, Kim M, Alexander T, Touret N, Hahn KM, Grinstein S. 2010. Amiloride inhibits macropinocytosis by lowering submembranous pH and preventing Rac1 and Cdc42 signaling. J Cell Biol 188:547–63.

70. Gao Y, Dickerson JB, Guo F, Zheng J, Zheng Y. 2004. Rational design and characterization of a Rac GTPase-specific small molecule inhibitor. Proc Natl Acad Sci U S A 101:7618–23.

71. Hansen CG, Nichols BJ. 2009. Molecular mechanisms of clathrin-independent endocytosis. J Cell Sci 122:1713–21.

72. Cureton DK, Massol RH, Whelan SP, Kirchhausen T. 2010. The length of vesicular stomatitis virus particles dictates a need for actin assembly during clathrin-dependent endocytosis. PLoS Pathog 6:e1001127.

73. Spearman P. 2018. Viral interactions with host cell Rab GTPases. Small GTPases 9:192–201.

74. Sieczkarski SB, Whittaker GR. 2003. Differential requirements of Rab5 and Rab7 for endocytosis of influenza and other enveloped viruses. Traffic 4:333–43.

75. Langemeyer L, Frohlich F, Ungermann C. 2018. Rab GTPase Function in Endosome and Lysosome Biogenesis. Trends Cell Biol 28:957–970.

76. Bohdanowicz M, Balkin DM, De Camilli P, Grinstein S. 2012. Recruitment of OCRL and Inpp5B to phagosomes by Rab5 and APPL1 depletes phosphoinositides and attenuates Akt signaling. Mol Biol Cell 23:176–87.

77. Choudhury A, Dominguez M, Puri V, Sharma DK, Narita K, Wheatley CL, Marks DL, Pagano RE. 2002. Rab proteins mediate Golgi transport of caveola-internalized glycosphingolipids and correct lipid trafficking in Niemann-Pick C cells. J Clin Invest 109:1541–50.

78. Tebaldi G, Pritchard, S.M., Nicola, A.V. 2020. Herpes simplex virus entry by a non-conventional endocytic pathway. J Virol doi:10.1128/JVI.01910-20.

79. Van Acker T, Tavernier J, Peelman F. 2019. The Small GTPase Arf6: An Overview of Its Mechanisms of Action and of Its Role in Host(-)Pathogen Interactions and Innate Immunity. Int J Mol Sci 20.

80. Garcia-Exposito L, Barroso-Gonzalez J, Puigdomenech I, Machado JD, Blanco J, Valenzuela-Fernandez A. 2011. HIV-1 requires Arf6-mediated membrane dynamics to efficiently enter and infect T lymphocytes. Mol Biol Cell 22:1148–66.

81. Heikkila O, Susi P, Tevaluoto T, Harma H, Marjomaki V, Hyypia T, Kiljunen S. 2010. Internalization of coxsackievirus A9 is mediated by {beta}2-microglobulin, dynamin, and Arf6 but not by caveolin-1 or clathrin. J Virol 84:3666–81.

82. Mercer J, Helenius A. 2008. Vaccinia virus uses macropinocytosis and apoptotic mimicry to enter host cells. Science 320:531–5.

83. Yoo JH, Shi DS, Grossmann AH, Sorensen LK, Tong Z, Mleynek TM, Rogers A, Zhu W, Richards JR, Winter JM, Zhu J, Dunn C, Bajji A, Shenderovich M, Mueller AL, Woodman SE, Harbour JW, Thomas KR, Odelberg SJ, Ostanin K, Li DY. 2016. ARF6 Is an Actionable Node that Orchestrates Oncogenic GNAQ Signaling in Uveal Melanoma. Cancer Cell 29:889–904.

84. Jackson CL, Walch L, Verbavatz JM. 2016. Lipids and Their Trafficking: An Integral Part of Cellular Organization. Dev Cell 39:139–153.

85. Musarrat F, Jambunathan N, Rider PJF, Chouljenko VN, Kousoulas KG. 2018. The Amino Terminus of Herpes Simplex Virus 1 Glycoprotein K (gK) Is Required for gB Binding to Akt, Release of Intracellular Calcium, and Fusion of the Viral Envelope with Plasma Membranes. J Virol 92.

86. Foster TP, Rybachuk GV, Kousoulas KG. 2001. Glycoprotein K specified by herpes simplex virus type 1 is expressed on virions as a Golgi complex-dependent glycosylated species and functions in virion entry. J Virol 75:12431–8.

87. Tal-Singer R, Peng C, Ponce De Leon M, Abrams WR, Banfield BW, Tufaro F, Cohen GH, Eisenberg RJ. 1995. Interaction of herpes simplex virus glycoprotein gC with mammalian cell surface molecules. J Virol 69:4471–83.

88. Komala Sari T, Gianopulos KA, Weed DJ, Schneider SM, Pritchard SM, Nicola AV. 2020. Herpes Simplex Virus Glycoprotein C Regulates Low-pH Entry. mSphere 5.

89. Rahn E, Petermann P, Hsu MJ, Rixon FJ, Knebel-Morsdorf D. 2011. Entry pathways of herpes simplex virus type 1 into human keratinocytes are dynamin- and cholesterol-dependent. PLoS One 6:e25464.

90. Clement C, Tiwari V, Scanlan PM, Valyi-Nagy T, Yue BY, Shukla D. 2006. A novel role for phagocytosis-like uptake in herpes simplex virus entry. J Cell Biol 174:1009–21.

91. Devadas D, Koithan T, Diestel R, Prank U, Sodeik B, Dohner K. 2014. Herpes simplex virus internalization into epithelial cells requires Na+/H+ exchangers and p21-activated kinases but neither clathrin-nor caveolin-mediated endocytosis. J Virol 88:13378–95.

92. Praena B, Bello-Morales R, Lopez-Guerrero JA. 2020. Hsv-1 Endocytic Entry into a Human Oligodendrocytic Cell Line is Mediated by Clathrin and Dynamin but Not Caveolin. Viruses 12.

93. Bid HK, Roberts RD, Manchanda PK, Houghton PJ. 2013. RAC1: an emerging therapeutic option for targeting cancer angiogenesis and metastasis. Mol Cancer Ther 12:1925–34.

94. Fretz M, Jin J, Conibere R, Penning NA, Al-Taei S, Storm G, Futaki S, Takeuchi T, Nakase I, Jones AT. 2006. Effects of Na+/H+ exchanger inhibitors on subcellular localisation of endocytic organelles and intracellular dynamics of protein transduction domains HIV-TAT peptide and octaarginine. J Control Release 116:247–54.

95. Gekle M, Drumm K, Mildenberger S, Freudinger R, Gassner B, Silbernagl S. 1999. Inhibition of Na+-H+ exchange impairs receptor-mediated albumin endocytosis in renal proximal tubule-derived epithelial cells from opossum. J Physiol 520 Pt 3:709–21.

96. Margiotta A, Bucci C. 2019. Coordination between Rac1 and Rab Proteins: Functional Implications in Health and Disease. Cells 8.

97. Huotari J, Helenius A. 2011. Endosome maturation. EMBO J 30:3481–500.

98. Gianni T, Campadelli-Fiume G, Menotti L. 2004. Entry of herpes simplex virus mediated by chimeric forms of nectin1 retargeted to endosomes or to lipid rafts occurs through acidic endosomes. J Virol 78:12268–76.

99. Zhang J, Feng Y, Forgac M. 1994. Proton conduction and bafilomycin binding by the V0 domain of the coated vesicle V-ATPase. J Biol Chem 269:23518–23.

100. Cheshenko N, Pierce C, Herold BC. 2018. Herpes simplex viruses activate phospholipid scramblase to redistribute phosphatidylserines and Akt to the outer leaflet of the plasma membrane and promote viral entry. PLoS Pathog 14:e1006766.

101. Zheng K, Chen M, Xiang Y, Ma K, Jin F, Wang X, Wang X, Wang S, Wang Y. 2014. Inhibition of herpes simplex virus type 1 entry by chloride channel inhibitors tamoxifen and NPPB. Biochem Biophys Res Commun 446:990–6.

102. Scott CC, Vacca F, Gruenberg J. 2014. Endosome maturation, transport and functions. Semin Cell Dev Biol 31:2–10.

103. Krummenacher C, Baribaud I, Sanzo JF, Cohen GH, Eisenberg RJ. 2002. Effects of herpes simplex virus on structure and function of nectin-1/HveC. J Virol 76:2424–33.

104. Stults AM, Smith GA. 2019. The Herpes Simplex Virus 1 Deamidase Enhances Propagation but Is Dispensable for Retrograde Axonal Transport into the Nervous System. J Virol 93.

105. Marconi P, Manservigi R. 2014. Herpes simplex virus growth, preparation, and assay. Methods Mol Biol 1144:19–29.

106. Dai X, Zhou ZH. 2014. Purification of Herpesvirus Virions and Capsids. Bio Protoc 4.

107. Kim DS, Dastidar H, Zhang C, Zemp FJ, Lau K, Ernst M, Rakic A, Sikdar S, Rajwani J, Naumenko V, Balce DR, Ewanchuk BW, Tailor P, Yates RM, Jenne C, Gafuik C, Mahoney DJ. 2017. Smac mimetics and oncolytic viruses synergize in driving anticancer T-cell responses through complementary mechanisms. Nat Commun 8:344.

108. Beug ST, Beauregard CE, Healy C, Sanda T, St-Jean M, Chabot J, Walker DE, Mohan A, Earl N, Lun X, Senger DL, Robbins SM, Staeheli P, Forsyth PA, Alain T, LaCasse EC, Korneluk RG. 2017. Smac mimetics synergize with immune checkpoint inhibitors to promote tumour immunity against glioblastoma. Nat Commun 8.

109. Li L, Wan T, Wan M, Liu B, Cheng R, Zhang R. 2015. The effect of the size of fluorescent dextran on its endocytic pathway. Cell Biol Int 39:531–9.

110. Schindelin J, Arganda-Carreras I, Frise E, Kaynig V, Longair M, Pietzsch T, Preibisch S, Rueden C, Saalfeld S, Schmid B, Tinevez JY, White DJ, Hartenstein V, Eliceiri K, Tomancak P, Cardona A. 2012. Fiji: an open-source platform for biological-image analysis. Nat Methods 9:676–82.

