## Supplemental Figures S1-S9 and Supplemental Table S1 for "Contributions of the four essential entry glycoproteins to HSV-1 tropism and the selection of entry routes"

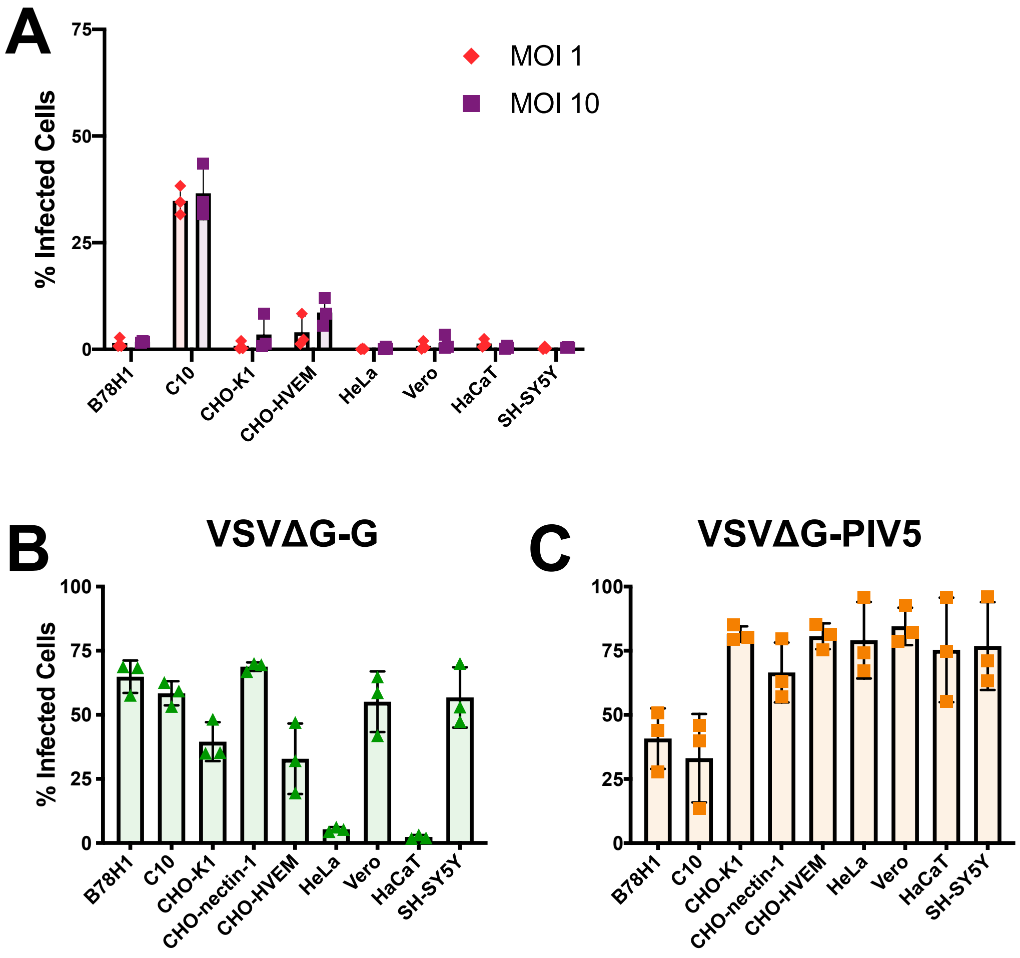


**Fig S1. Infecting cells with VSVΔG-BHLD at a higher MOI does not increase entry to an appreciable extent.** A)Receptor null (B78H1 and CHO-K1) and receptor bearing cells (C10, CHO-HVEM, HeLa, Vero, HaCaT, and SH-SY5Y) were infected at MOI =1 (red) or MOI = 10 (purple). Entry efficiency was assessed by flow cytometry at 6 hours post infection. B and C) Receptor null (B78H1 and CHO-K1) and receptor bearing cells (C10, CHO-HVEM, HeLa, Vero, HaCaT, and SH-SY5Y) were infected at MOI =1 with either VSVΔG-G (B) or VSVΔG-PIV5 (C). Entry was assessed by flow cytometry at 6 hours post infection.

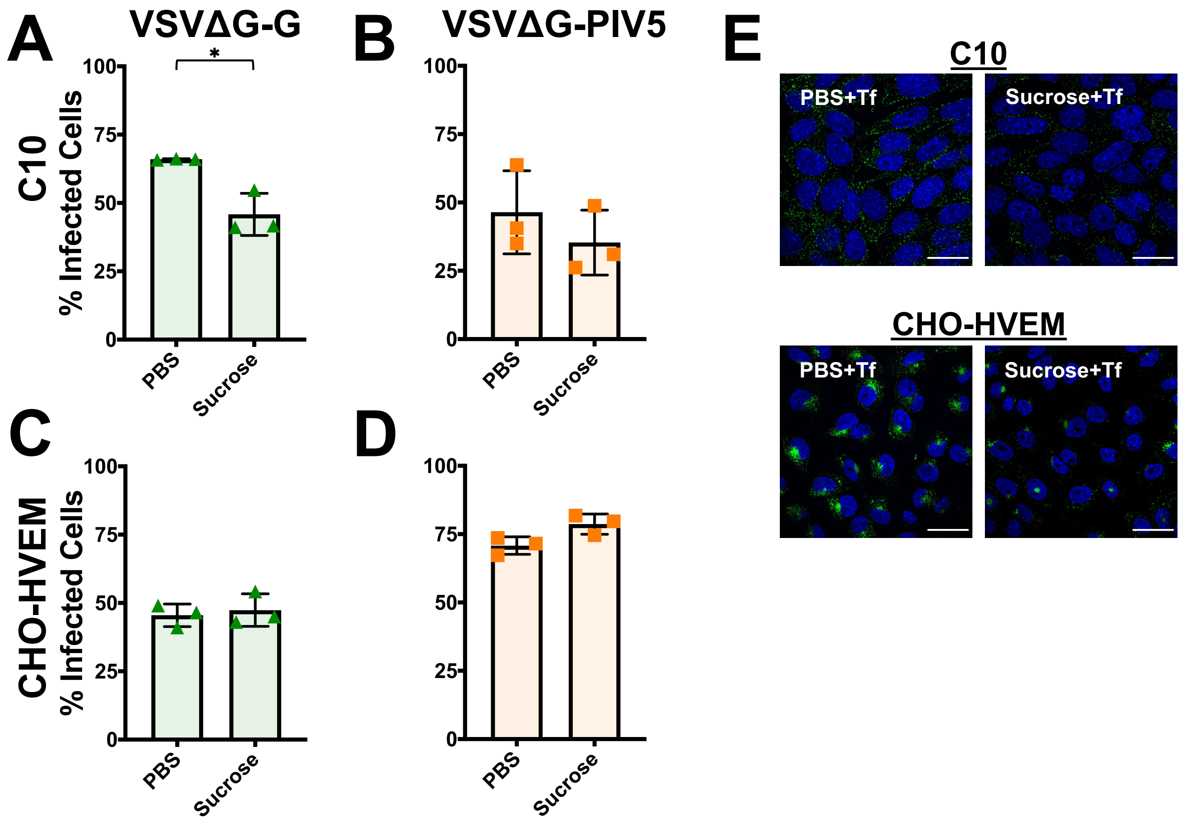


**Fig. S2. VSVΔG-G and VSVΔG-PIV5 entry in the presence of hypertonic sucrose.** C10 (A, B) and CHO-HVEM (C, D) cells were pretreated with a hypertonic solution of sucrose (0.3 M) and infected with VSVΔG-G or VSVΔG-PIV5 at MOI = 1. Infectivity was quantitated by flow cytometry at 6 hours post infection. Significance was calculated using a two-tailed Student's T-test with Welch's correction ($p < 0.05$ = *; $p < 0.01$ = **; $p < 0.001$ = ***). E) C10 and CHO-HVEM cells were pretreated with 0.3 M sucrose and incubated with 50 ug/ml of AF488-labeled transferrin (Tf). Cells were fixed, counterstained with DAPI, and imaged by confocal microscopy. Scale bar = 25 $\mu$m.

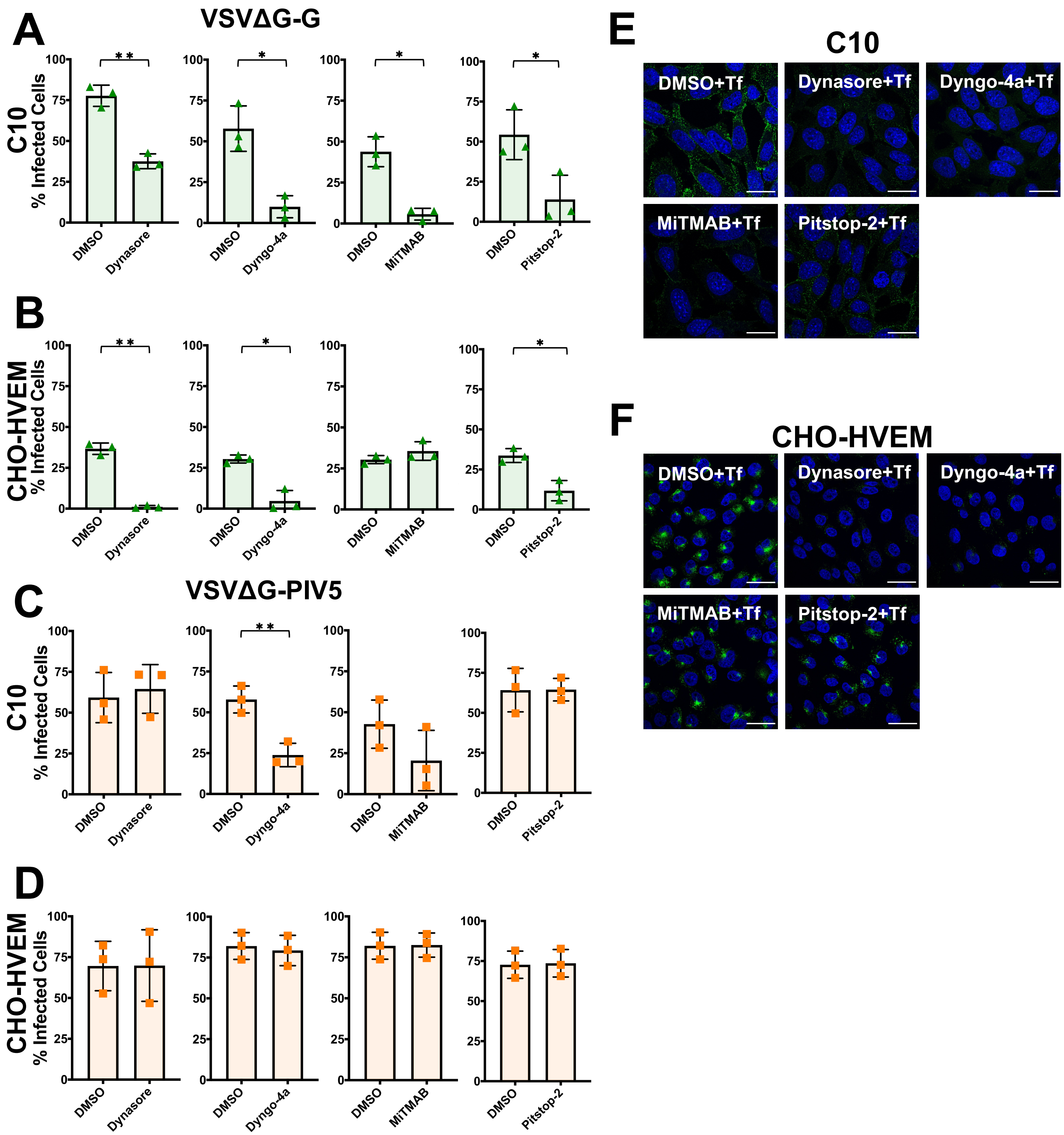


**Fig. S3 VSVΔG-G and VSVΔG-PIV5 differ in their dependence on dynamin and clathrin for entry.** C10 (A and C) and CHO-HVEM (B and D) cells were pretreated with dynamin inhibitors Dynasore (80 μM), Dyngo-4a (25 μM), MiTMAB (5 μM), or the CME inhibitor Pitstop-2 (30 μM) and infected with VSVΔG-G or VSVΔG-PIV5 at a MOI of 1. Infectivity was quantitated by flow cytometry at 6 hours post infection. CHO-HVEM cells treated with Dyngo-4a or MiTMAB used the same DMSO control as indicated by the same bar graph appearing twice each in panels C and D. Significance was calculated using a two-tailed Student's T-test with Welch's correction ($p < 0.05$ = *; $p < 0.01$ = **; $p < 0.001$ = ***). E and F) C10 and CHO-HVEM cells were pretreated with dynamin inhibitors Dynasore, Dyngo-4a, MiTMAB, or Pitstop-2 at the same concentrations as in panels A-D and then incubated with 50 μg/ml of AF488-labeled transferrin. Cells were fixed, counterstained with DAPI, and imaged by confocal microscopy. Scale bar = 25 μm.

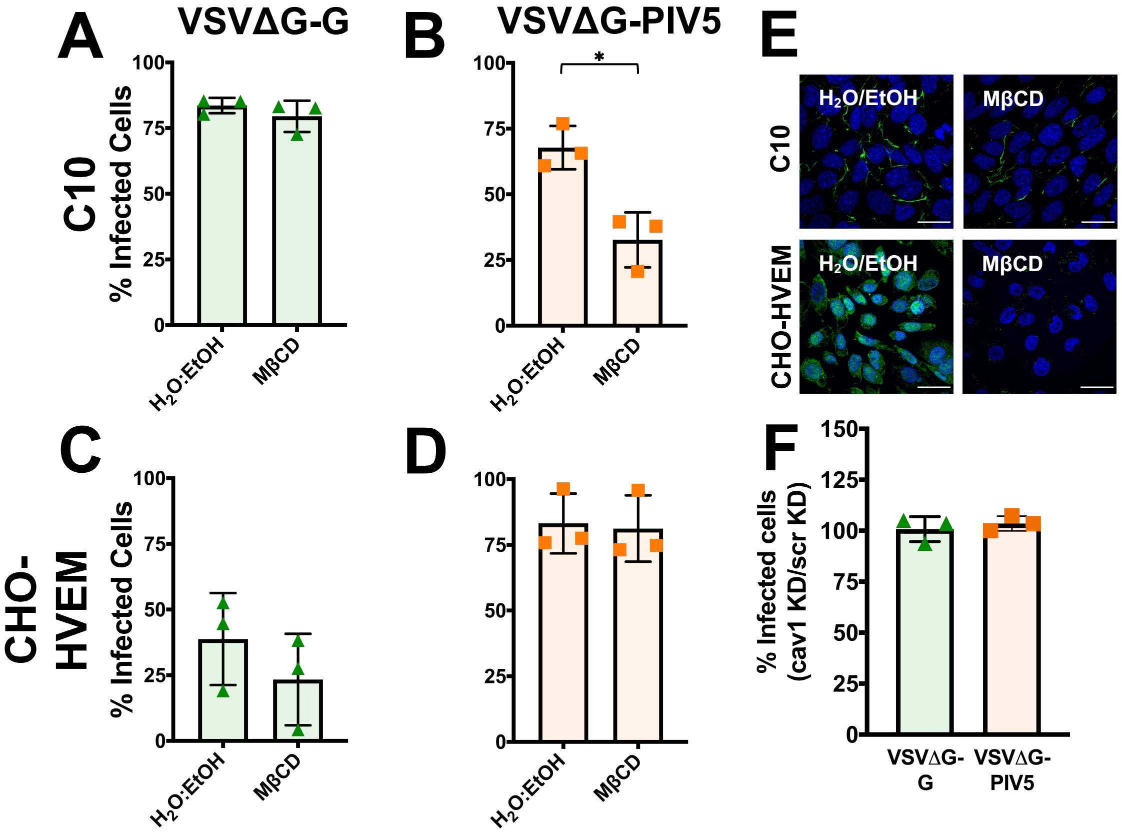


**Fig. S4. VSVΔG-G entry does not require cholesterol whereas VSVΔG-PIV5 entry requires cholesterol in a cell-type-dependent manner.** C10 (A and B) and CHO-HVEM (C and D) cells were pretreated with a cholesterol-removal drug methyl-β-cyclodextran, MβCD (5 mM) and infected with VSVΔG-G or VSVΔG-PIV5 at a MOI of 1. Infectivity was quantitated by flow cytometry at 6 hours post infection. E) C10 and CHO-HVEM cells were treated with either a solvent control ($H_2O$/EtOH) or methyl-β-cyclodextrin (MβCD), then incubated with cholera toxin subunit B labelled with Alexa Fluor 488. Confocal microscopy was performed on the solvent control and methyl-β-cyclodextrin treated cells. Cells were fixed, counterstained with DAPI, and imaged by confocal microscopy. Scale bar = 25 μm. (F) CHO-HVEM cells were transfected with a caveolin-1 siRNA (cav-1) or a scrambled control siRNA (scr) (both 50 pm) and infected with VSVΔG-G or VSVΔG-PIV5 at a MOI of 1. Infectivity was quantitated by flow cytometry at 6 hours post infection. Significance was calculated using a two-tailed Student's T-test with Welch's correction ($p < 0.05$ = *; $p < 0.01$ = **; $p < 0.001$ = ***).

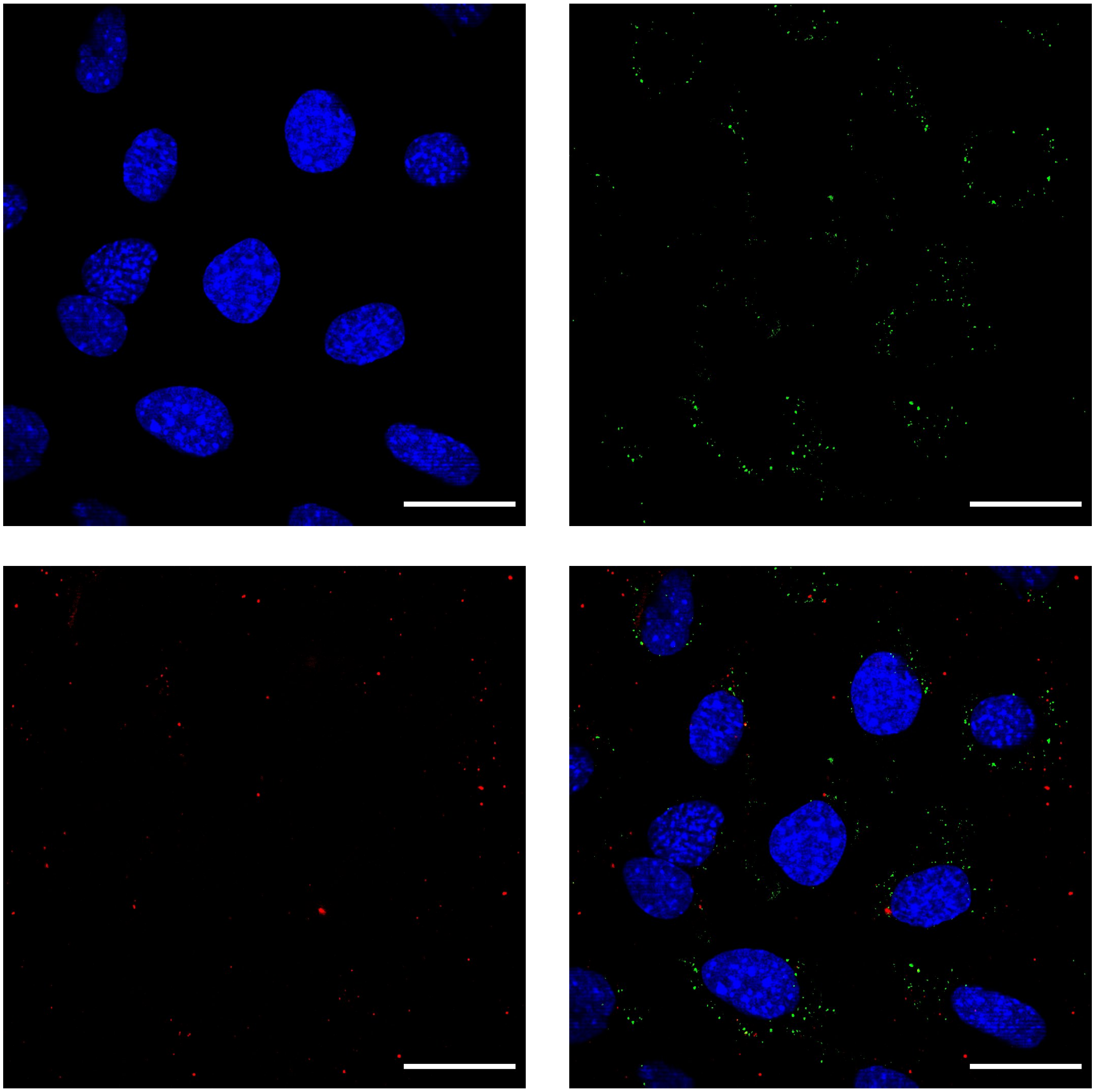

**Fig. S5. VSVΔG-BHLD does not co-localize with the fluid-phase marker 70 kDa dextran in C10 cells.** C10 cells were incubated with 1 mg/ml of rhodamine-B labelled 70 kDa dextran and VSVΔG-BHLD (MOI = 1) for one hour at 4˚C. Cells were then shifted to 37 ˚C for 20 minutes. Cells were fixed, counterstained with DAPI, and imaged by confocal microscopy. gB was detected by immunofluorescence using the rabbit pAb R68 and anti-rabbit IgG conjugated to FITC. Green = gB (marker for VSVΔG-BHLD particles); Red = 70 kDa dextran. Scale bar = 25 μm.

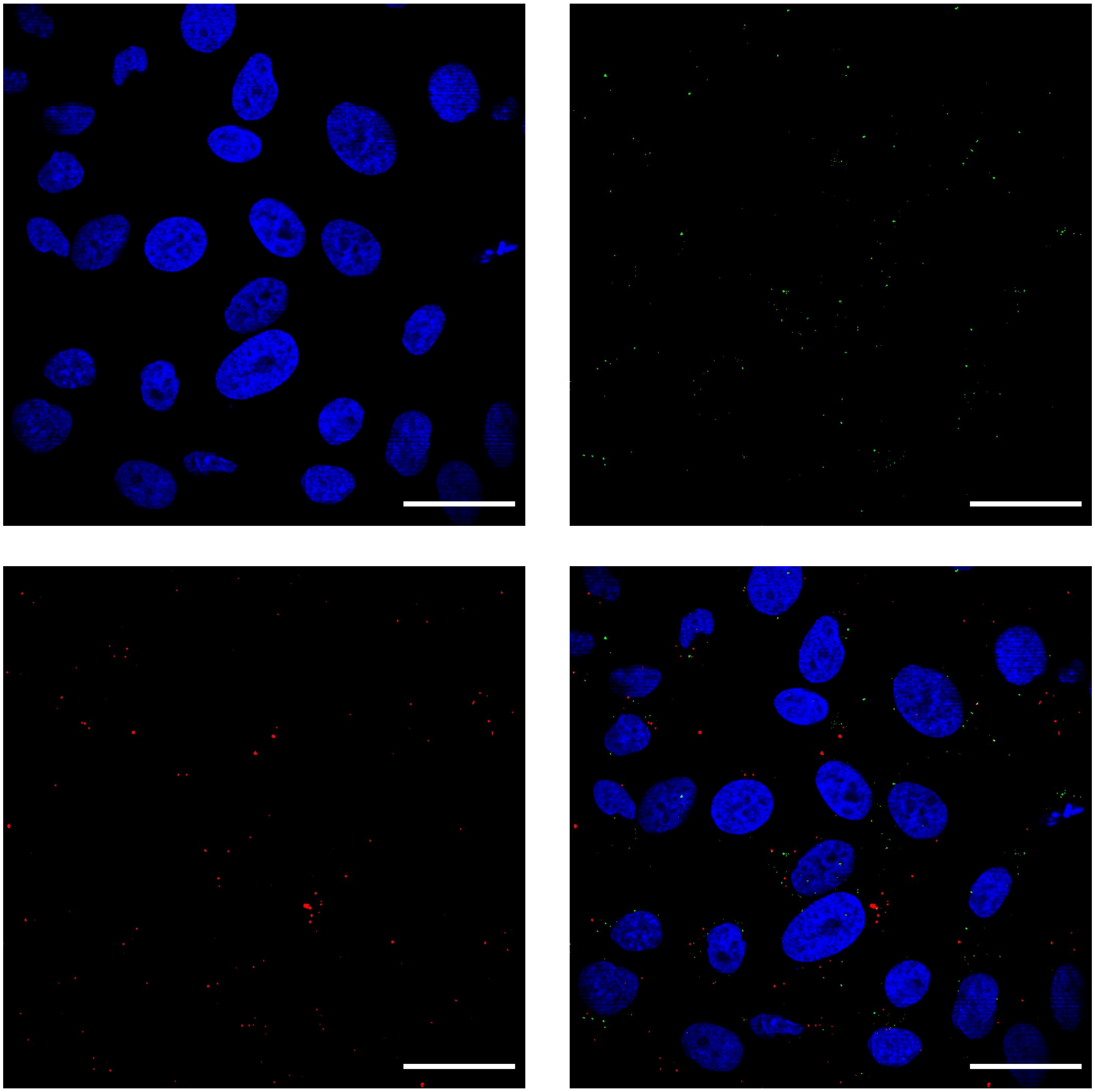

**Fig. S6. VSVΔG-BHLD, in large, does not co-localized with the fluid-phase marker, 70 kDa dextran, in CHO-HVEM cells.** CHO-HVEM cells were incubated with 1 mg/ml of rhodamine-B labelled 70 kDa dextran and VSVΔG-BHLD (MOI = 1) for one hour at 4˚C. Cells were then shifted to 37˚C for 20 minutes. Cells were fixed, counterstained with DAPI, and imaged by confocal microscopy. gB was detected by immunofluorescence using the rabbit pAb R68 and anti-rabbit IgG conjugated to FITC. Green = gB (marker for VSVΔG-BHLD particles); Red = 70 kDa dextran. Scale bar = 25 μm.

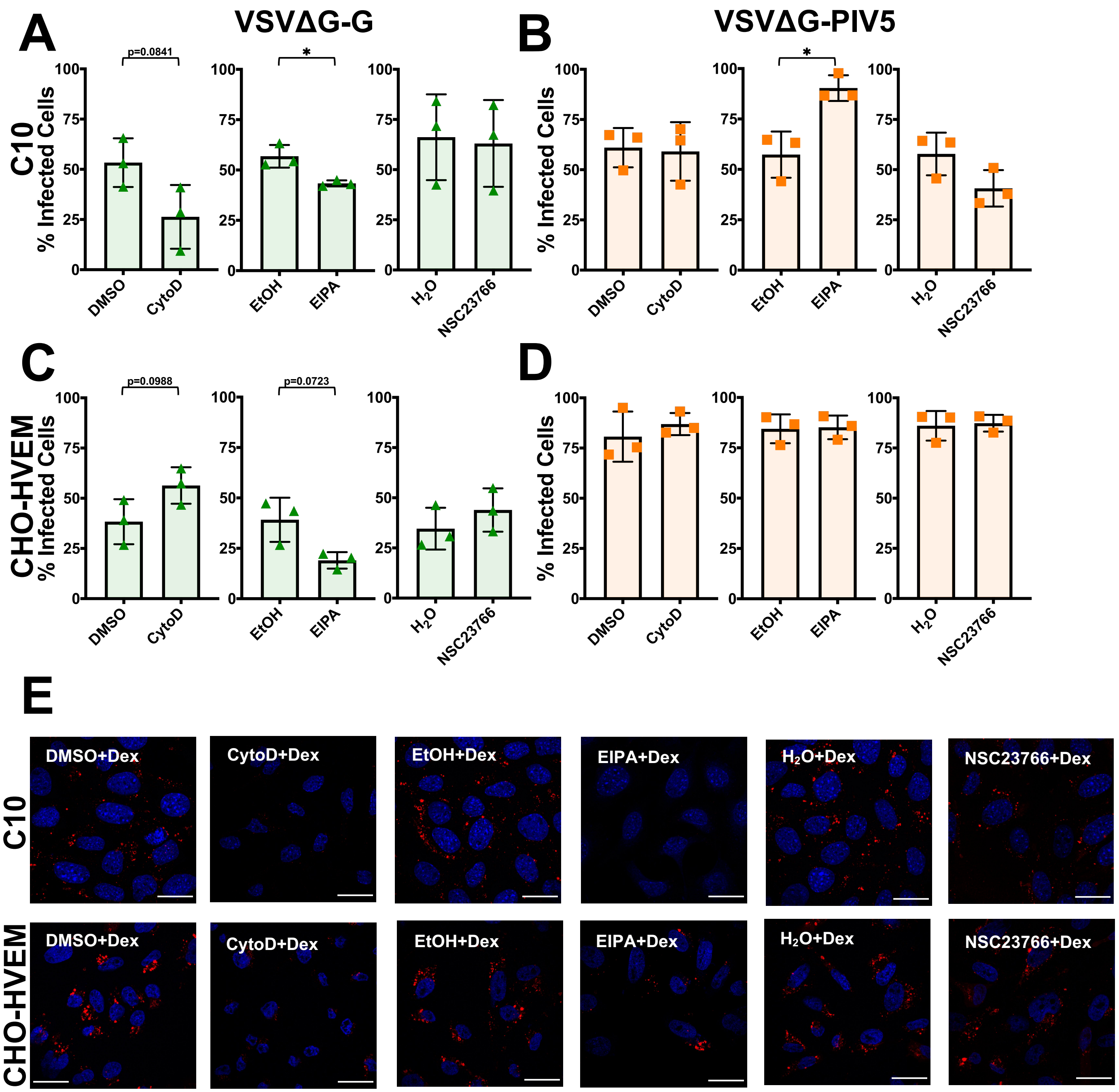


**Fig. S7. VSVΔG-G and VSVΔG-PIV5 entry does not require macropinocytosis.** C10 (A and B) and CHO-HVEM (C and D) cells were pretreated with macropinocytosis inhibitors cytochalasin D (2 μM), EIPA (25 μM), or NSC23766 (200 μM) and infected with VSVΔG-G or VSVΔG-PIV5 at a MOI of 1. Infectivity was quantitated by flow cytometry at 6 hours post infection. Significance was calculated using a two-tailed Student's T-test with Welch's correction ($p < 0.05$ = *; $p < 0.01$ = **; $p < 0.001$ = ***). E) C10 and CHO-HVEM cells were pretreated with macropinocytosis inhibitors cytochalasin D, EIPA, or NSC23766 at the same concentrations as in panels A-D and then incubated with 1.0 mg/ml of Rhodamine-B-labeled 70-kDa dextran (Dex). Cells were fixed, counterstained with DAPI, and imaged by confocal microscopy. Scale bar = 25 μm.

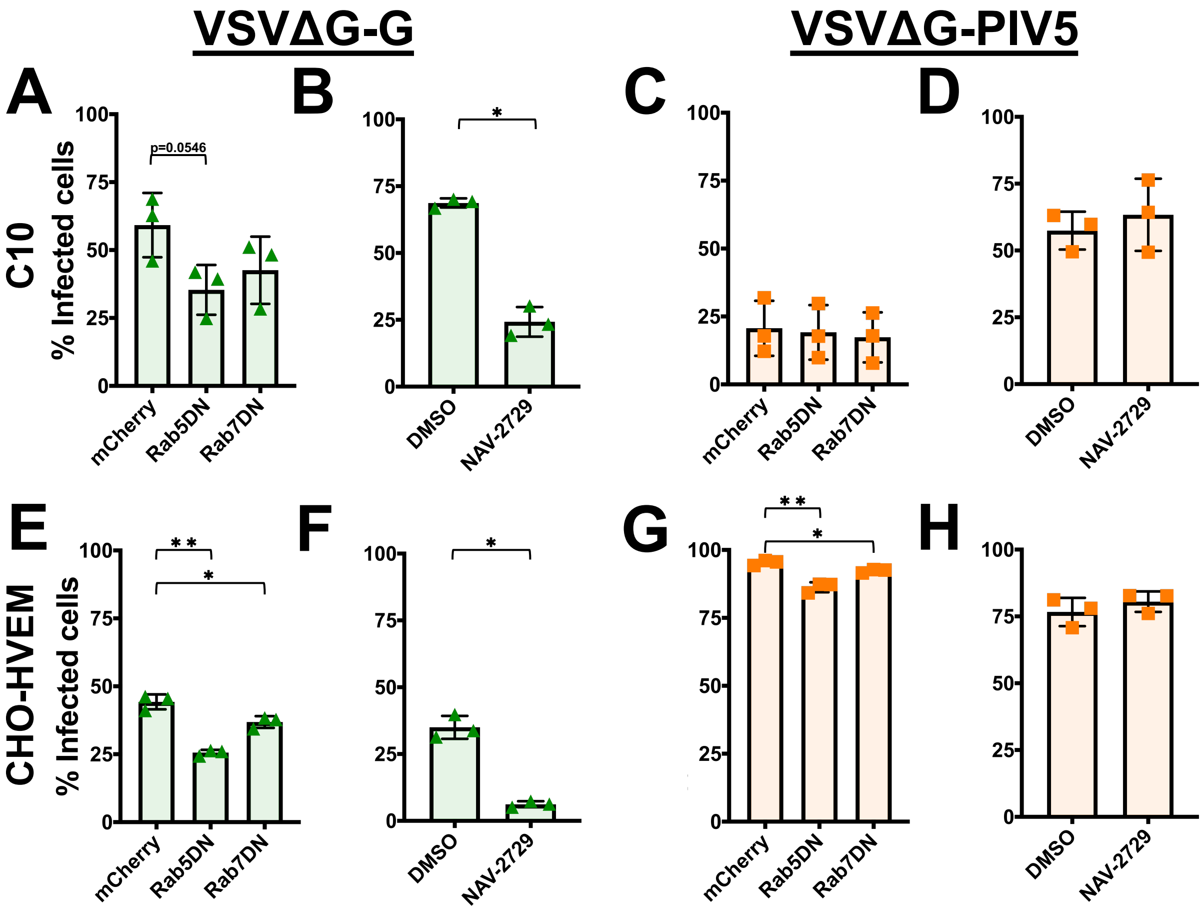


**Fig. S8**: **Roles of Rab5, Rab7, and Arf6 in VSVΔG-G and VSVΔG-PIV5 entry.** The roles of the small GTPases Rab5, Rab7 (A, C, E, G), and Arf6 (B, D, F, H) were assessed for VSVΔG-G (A, B, E, F) and VSVΔG-PIV5 (C, D, G, H) entry into C10 (A, B, C, D) and CHO-HVEM (E, F, G, H) cells. C10 (A and C) and CHO-HVEM (E and G) cells were transfected with either an empty vector control (eGFP or mCherry), eGFP or mCherry-tagged Rab5 dominant negative (DN), or eGFP or mCherry-tagged Rab7DN. Cells were infected at an MOI = 1 with either VSVΔG-G or VSVΔG-PIV5. Entry was assessed by flow cytometry at 6 hpi. The percent of infected cells was determined by dividing the number of virus(+)/eGFP/mCherry(+) cells by the total number of eGFP/mCherry(+) cells. C10 (B and D) and CHO-HVEM cells (F and H) were treated with the Arf6 inhibitor NAV-2729 (25 μM) and infected with either VSVΔG-G or VSVΔG-PIV5 at an MOI = 1. Significance was calculated using a two-tailed Student's T-test with Welch's correction (ns = not significant; $p < 0.05$ = *; $p < 0.01$ = **; $p < 0.001$ = ***).

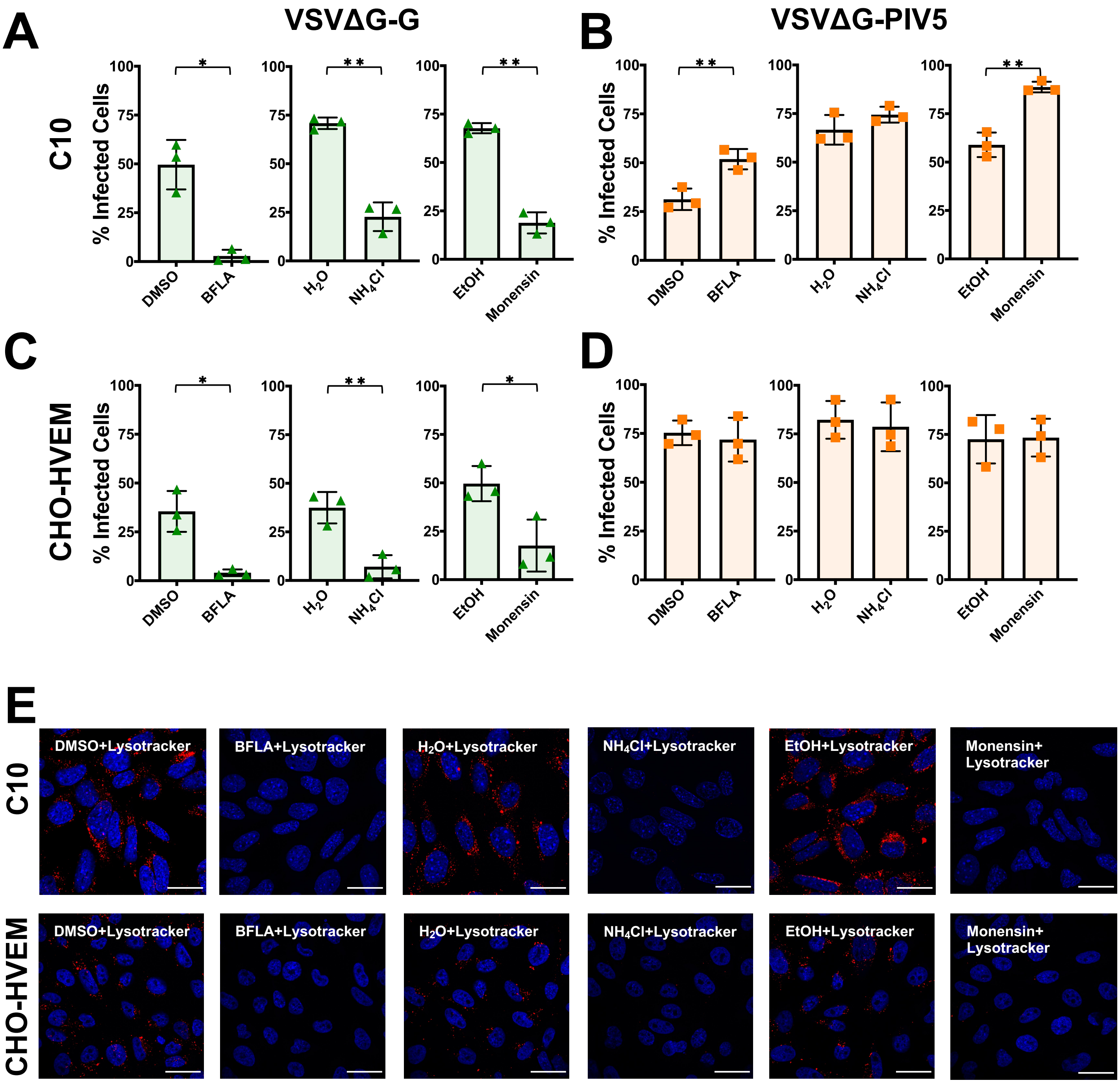


**Fig. S9. VSVΔG-G but not VSVΔG-PIV5 entry requires endosomal acidification.** C10 (A and B) and CHO-HVEM (C and D) cells were pretreated with inhibitors of endosomal acidification BFLA (100 nM), $NH_4Cl$ (50 mM), or monensin (15 μM) and infected with VSVΔG-G or VSVΔG-PIV5 at MOI = 1. Infectivity was quantitated by flow cytometry at 6 hours post infection. Significance was calculated using a two-tailed Student's T-test with Welch's correction ($p < 0.05$ = *; $p < 0.01$ = **; $p < 0.001$ = ***). E) C10 and CHO-HVEM cells were pretreated with inhibitors of endosomal acidification at the same concentrations as in panels A-D (BFLA, $NH_4Cl$, or monensin) and then incubated with Lysotracker (1 μM). Cells were fixed, counterstained with DAPI, and imaged by confocal microscopy. Scale bar = 25 μm.

| Target | Inhibitors | HSV-1 | | VSVΔG-BHLD | | VSVΔG-G | | VSVΔG-PIV5 | |
|---|---|---|---|---|---|---|---|---|---|
| | | C10 | CHO-HVEM | C10 | CHO-HVEM | C10 | CHO-HVEM | C10 | CHO-HVEM |
| Endocytosis | Hypertonic sucrose | ✓ | ✓ | ✓ | ✓ | ✓ | X | X | X |
| Dynamin | Dynasore | ✓ | ✓ | ✓ | ✓ | ✓ | ✓ | X | X |
| | Dyngo-4a | ✓ | ✓ | ✓ | ✓ | ✓ | ✓ | ✓ | X |
| | MiTMAB | ✓ | ✓ | ✓ | ✓ | ✓ | ✓ | X | X |
| Clathrin | Pitstop-2 | ✓ | ✓ | X | X | ✓ | ✓ | X | X |
| Cholesterol | MβCD | ✓ | ✓ | ✓ | ✓ | X | X | ✓ | X |
| Caveolin | Cav-1 siRNA | X | X | X | X | X | X | X | X |
| Macropinocytosis | CytoD | X | X | X | X | ✓ | X | X | X |
| | EIPA | X | ✓ | X | X | ✓ | ✓ | X | X |
| | NSC23766 | X | X | ✓ | ✓ | X | X | X | X |
| Small GTPases | Rab5DN | X | X | ✓ | X | ✓ | ✓ | X | X |
| | Rab7DN | X | X | X | X | X | X | X | X |
| | NAV-2729 | ✓ | ✓ | X | X | ✓ | ✓ | X | X |
| Low pH | BFLA | X | X | ✓ | X | ✓ | ✓ | X | X |
| | $NH_4Cl$ | ✓ | ✓ | ✓ | ✓ | ✓ | ✓ | X | X |
| | Monensin | ✓ | ✓ | ✓ | X | ✓ | ✓ | X | X |

**Table S1. Sensitivity of HSV-1, VSVΔG-BHLD, VSVΔG-G, and VSVΔG-PIV5 to specific inhibitors.** Green check marks indicate that virus entry is sensitive to that particular inhibitor. Red X marks indicate that the virus is not sensitive to that particular inhibitor.
